# mTOR inhibition in Q175 Huntington’s disease model mice facilitates neuronal autophagy and mutant huntingtin clearance

**DOI:** 10.1101/2024.05.29.596471

**Authors:** Philip Stavrides, Chris N. Goulbourne, James Peddy, Chunfeng Huo, Mala Rao, Vinod Khetarpal, Deanna M. Marchionini, Ralph A. Nixon, Dun-Sheng Yang

## Abstract

Huntington’s disease (HD) is caused by expansion of the polyglutamine stretch in huntingtin protein (HTT) resulting in hallmark aggresomes/inclusion bodies (IBs) composed of mutant huntingtin protein (mHTT) and its fragments. Stimulating autophagy to enhance mHTT clearance is considered a potential therapeutic strategy for HD. Our recent evaluation of the autophagic-lysosomal pathway (ALP) in human HD brain reveals upregulated lysosomal biogenesis and relatively normal autophagy flux in early Vonsattel grade brains, but impaired autolysosome clearance in late grade brains, suggesting that autophagy stimulation could have therapeutic benefits as an earlier clinical intervention. Here, we tested this hypothesis by crossing the Q175 HD knock-in model with our autophagy reporter mouse TRGL (**T**hy-1-**R**FP-**G**FP-**L**C3) to investigate *in vivo* neuronal ALP dynamics. In the Q175 and/or TRGL/Q175 mice, mHTT was detected in autophagic vacuoles and also exhibited a high level of colocalization with autophagy receptors p62/SQSTM1 and ubiquitin in the IBs. Compared to the robust lysosomal pathology in late-stage human HD striatum, ALP alterations in Q175 models are also late-onset but milder that included a lowered phospho-p70S6K level, lysosome depletion and autolysosome elevation including more poorly acidified autolysosomes and larger-sized lipofuscin granules, reflecting impaired autophagic flux. Administration of a mTOR inhibitor to 6-mo-old TRGL/Q175 normalized lysosome number, ameliorated aggresome pathology while reducing mHTT-, p62- and ubiquitin-immunoreactivities, suggesting the beneficial potential of autophagy modulation at early stages of disease progression.

## Introduction

Huntington’s disease (HD) is an autosomal dominant disorder caused by a mutation in the gene encoding the huntingtin protein (HTT) resulting in expansion of the polyglutamine (polyQ) stretch on its amino-terminus [7, 18, 26, 71]. HD pathogenesis advances in a spatiotemporal pattern, which is used to stage disease pathology severity as Grade 0-4 (HD0, HD1, HD2, etc.)[76]. GABA-containing medium spiny projection neurons in the striatum are most susceptible to cell death [75], and the cerebral cortex, particularly layer 5a, also shows cell loss [57, 68]. Neuronal intranuclear inclusions (NIIs) and neuropil inclusions are present in HD brains and are positive for mutant huntingtin (mHTT) and ubiquitin (Ub) [17, 20].

Recently, we completed a comprehensive evaluation of the autophagic-lysosomal pathway (ALP) in the HD brain at progressive disease stages, focusing on the most affected brain region, the striatum, in comparison to a less affected region, the neocortex [9]. Double fluorescence immunolabeling and immuno-electron microscopy (IEM) revealed colocalization of HTT/mHTT with the autophagy-related adaptor proteins, p62/SQSTM1 and ubiquitin, and cathepsin D (CTSD) within aggresome inclusions and autophagic compartments, documenting the involvement of ALP in HTT/mHTT turnover and the disease-related impairment of this process in late-stage disease. The temporal evolution of ALP alterations generally revealed minimal detectable impairment of upstream autophagy steps [e.g., autophagosome (AP) induction, formation, and fusion with lysosomes (LY)] and, in striatum, elevated levels of LAMP1 and LAMP2 markers suggesting modestly upregulated LY biogenesis. At late disease stages, mainly HD4, neuronal ALP dysfunction exhibited enlarged/clumped CTSD-immunoreactive autolysosomes (AL)/LY and ultrastructural evidence of autophagic vacuole (AV) fusion and transition to lipofuscin granule formation. These findings collectively suggest that relatively competent autophagy machinery is maintained during the disease progression with a compensatory upregulation in lysosomal biogenesis, which together prevents against mHTT accumulation. This situation is failing at the late disease stages when AL clearance is impeded, substrates, including mHTT and its metabolites, accumulate in AL and aggresome inclusions increase.

A possible implication of the foregoing findings is that pharmacologic enhancement of autophagy applied at a symptomatic but early stage of disease, when the ALP clearance machinery is fully competent, may be therapeutic in clearing mHTT protein in affected neurons. By contrast, in Alzheimer’s Disease (AD) lysosomal clearance deficits develop at the earliest disease stages, suggesting enhanced autophagy induction is counterproductive.

Modulation of autophagy as a therapeutic strategy for HD has been investigated in various cell and animal models of HD [11, 26, 28, 66, 81]. Of particular relevance to our present study are those involving autophagy induction with mTOR-dependent or -independent autophagy-enhancing approaches in HD mouse models [e.g. Trehalose [70]; Rapamycin analog Temsirolimus (CCI-779) [59]; Rilmenidine [61]; Rhes manipulations [5, 36]]. These studies have generally demonstrated ameliorative effects on outcome measures such as mHTT lowering and behavioral/motor function assays. However, in many cases it is unclear whether autophagy mechanisms are directly engaged in the brain or are critical to rescue.

Thus, our study aimed to comprehensively characterize the autophagy response with a range of autophagy markers to interrogate the competence of the entire autophagy process in relationship to mHTT in the well-characterized zQ175 Knock-In HD mouse model (Q175). Towards this goal, we evaluated autophagy in neurons of Q175 after introducing by neuron-specific transgenesis the dual fluorescence-tagged autophagy probe, **t**andem **f**luorescent mRFP-eGFP-**LC3** (tfLC3), an autophagy adaptor protein associated with AP and degraded via autophagy [35]. We thereby generated a new Q175 cross, namely TRGL **(T**hy-1-**R**FP-**G**FP-**L**C3)/Q175. tfLC3 expression, driven postnatally by the neuron-specific Thy1-promoter, allows for selective monitoring of neuronal autophagy without the confounding influence of glial cells. Resolution and sensitivity in reporting tfLC3 signal is high compared to conventional immunofluorescence staining of LC3. The ability to ratiometrically report pH-dependent changes in fluorescence (hue angle) enables neutral pH AP to be distinguished from acidic AL, that progressively acidify intraluminally upon fusion with LY. We are further able to differentiate AL subgroups differing in their extent of acidification after LY fusion. Distinguishing properly acidified AL from those poorly acidified because of delayed or defective acidification is assisted by immunolabeling AL with a marker, such as CTSD, which is then detected with a third fluorophore. Together this triple fluorescence paradigm is objectively quantified by computer-assisted deconvolution of the proportions of each label within the analyzed neuron, a reflection of their relative pH (and fusion with LY), which allows ALP organelle subtypes including LC3-nagative LY to be identified and quantified for their numbers, sizes and spatial distributions in intact brain sections. The collective data provides reports on the completion (or lack thereof) of autophagy flux (ALP dynamics) and any blockage at particular steps in the ALP pathway, including the normal acidification and further maturation of AL through their successful elimination of fluorescence-tagged LC3 [35, 37, 39, 40].

Crossing TRGL mice with a model of a neurodegenerative disease, the Q175 mouse model of HD, has enabled us to assess disease-related autophagy alterations in the Q175 and TRGL/Q175 models and their response to a pharmacological inhibition of mTOR, (mTORi) INK-128 (hereafter INK). Our data demonstrate target engagement and positive effects of the compound on rescuing Q175 phenotypes including reversal on AL/LY subtypes as reported by the tfLC3 probe and parallel reductions of mHTT-, p62- and Ub-immunoreactivity (IR), suggesting that the compound targeted the ALP to degrade mHTT.

## Materials and Methods

### The HD Q175 mouse model and the generation of a new Q175 model crossed with the autophagy reporter mouse TRGL (TRGL/Q175)

Q175 mice (B6J.zQ175 Knock-In mice, CHDI- 81003003)[47, 48] were obtained from the Jackson Lab (Stock No. 027410) on a C57BL/6J background. The zQ175 KI allele has the mouse *Htt* exon 1 replaced by the human *HTT* exon 1 sequence containing a ∼190 CAG repeat tract, and the average size for the mice used in this study was 200 CAG. Q175 genotyping and CAG sizing were conducted by Laragen, Inc. (Culver City, CA). The TRGL (**T**hy-1 m**R**FP-e**G**FP-**L**C3) mouse model, expressing tandem fluorescence-tagged LC3 (tfLC3 or mRFP-eGFP-LC3) in neurons, was generated, genotyped by PCR and maintained in a C57BL/6J background as described previously [35]. The TRGL was crossed with Q175 to generate the new TRGL/Q175 model featuring the tfLC3 probe as an *in vivo* autophagy reporter.

The mice were maintained in the Nathan Kline Institute for Psychiatric Research (NKI) animal facility and housed in a 12 hrs light/dark cycle. All animal procedures were performed following the National Institutes of Health Guidelines for the Humane Treatment of Animals, with approval from the Institutional Animal Care and Use Committee at the NKI. Animals of both sexes were used in this study. Details for mouse ages are in the Figure Legend and/or Results. All efforts were made to minimize animal suffering and the number of animals used.

### mTORi INK administration to mice

INK-128 (ChemScene) was formulated in the sterile filtered vehicle (Veh) – 0.5% carboxymethyl cellulose (Sigma, Cat #5678) and 0.05% tween 80 in water – at different concentrations to achieve the various desired dosages. The mixture was homogenized with a tissue homogenizer, stored in a sterile container at room temperature for up to 2 weeks, and stirred prior to each dosing. It was administered to mice via oral gavage at a dose volume of 5 ml/kg, which was a relatively smaller volume of drug solution (generally 100-150 ul depending on the mouse body weight) aimed at avoiding the suppression of appetite. Mice receiving the Veh (also at a solution volume of 5 ml/kg) served as the control group. Oral gavage was conducted with 20G-38mm Cadence Science™ Malleable Stainless-Steel Animal Feeding Needles #9921 (Fisher Sci Cat #.14-825-275).

### mTORi INK pharmacokinetics in mouse brains after oral administration

INK was formulated into 0.5% carboxymethyl cellulose and 0.05% tween-80 in water to be administered orally. 6 months old C57B6/J male mice were treated with compound at 1, 3 or 10 mg/kg (mpk) or vehicle. 6, 12 or 24 hours after administration, mice were terminally anesthetized with pentobarbital and cerebellum was dissected and fresh frozen on liquid nitrogen and stored at -80°C until analysis. Tissue was homogenized and extracted with acetonitrile (containing 0.1% formic acid). Extract was analyzed using a LC-MS/MS (Waters Xevo TQ MS) method with a lower limit of quantitation of 13 nM.

### Brain tissue preparation

To obtain tissues for experiments, the animals were anesthetized with a mixture of ketamine (100 mg/kg BW) and xylazine (10 mg/kg BW). Mice for light microscopic (LM) analyses were usually fixed by cardiac perfusion using 4% paraformaldehyde (PFA) in 0.1 M sodium cacodylate buffer [pH 7.4, Electron Microscopy Sciences (EMS), Hatfield, PA]. Following perfusion fixation, the brains were immersion-fixed in the same fixative overnight at 4°C. For transmission electron microscopic (EM) study, 4% PFA was supplemented with 2% glutaraldehyde (EMS). For biochemical analyses such as immunoblotting, the brains were flash frozen on dry ice and stored at –70°C. When both morphological and biochemical analyses were to be performed on the same brain, the brain was removed after brief perfusion with saline. One hemisphere was frozen at –70°C and the other half was immersion-fixed in 4% PFA for 3 days at 4°C.

### Antibodies for immunohistochemistry (IHC) and western blotting (WB)

The following primary antibodies were used in this study. <u>(1) from Cell Signaling Technology:</u> tHTT rabbit mAb (total HTTs, #5656), MTOR rabbit mAb (#2983); p-MTOR (S2481 autophosphorylation) pAb (#2974), p-MTOR (S2448) pAb (#2971), p70S6K rabbit mAb (#2708), p-p70S6K (S371) pAb (#9208), p-p70S6K (T389) rabbit mAb (#9235), S6 ribosomal protein mAb (#2317), p-S6 ribosomal protein (S240/244) rabbit mAb (#5364), ULK1 pAb (#4773), ULK1 rabbit mAb (#6439), p-ULK1 (S757) pAb (#6888), p-ULK1 (S317) pAb (#37762), ATG5 rabbit mAb (#12994), ATG14 pAb (#5504), p-ATG14 (S29) rabbit mAb (#92340), p-Beclin 1 (S30) rabbit mAb (#35955), VPS34 rabbit mAb (#4263), TRAF6 rabbit mAb (#8028). <u>(2) from Millipore-Sigma:</u> ntHTT mAb (mEM48, #MAB5374; N-Terminus-specific), HTT mAb (1C2, #MAB1574; epitope: N-terminal part of the human TATA Box Binding Protein (TBP) containing a 38-glns stretch), HTT mAb (#MAB5490; epitope: human HTT aa115-129), ATG5 pAb (#ABC14), LC3 pAb (#ABC929), β−actin mAb (#A1978). <u>(3) from other vendors:</u> HTT mAb MW8 (epitope: HD exon 1 with 67Q; Develop Studies Hybridoma Bank, University of Iowa), HTT mAb PHP2 (CHDI-90001516-2, Coriell/CHDI), Beclin 1 mAb (BD Biosciences, #612113), LC3 pAb (Novus Biologics, #NB100- 2220), p62/SQSTM1 mAb (BD Biosciences, #610832) or C-term-specific p62/SQSTM1 Guinea Pig pAb (Progen Biotechnik #C-1620); ubiquitin pAb (Dako Agilent, #Z0458), ubiquitin pAb (Abcam, #ab7780), LAMP1 rat mAb (Developmental Studies Hybridoma Bank, #H4A3), CTSB goat pAb (Neuromics, #GT15047). <u>(4) In-house made:</u> CTSD pAb (RU4). CTSD sheep pAb (D- 2-3)[13].

The following secondary antibodies and reagents for immunoperoxidase labeling were purchased from Vector Laboratories (Burlingame, CA): biotinylated goat anti-rabbit or -mouse IgG/IgM, Vectastain ABC kit (PK-4000), and DAB Peroxidase Substrate Kit (SK-4100). Mouse on Mouse (M.O.M) detection kit (BMK-2201), normal-goat (S-100) and normal-donkey (S-2000- 20) serum blocking solution were also from Vector Lab. The following secondary antibodies for immunofluorescence were purchased from Thermo Fisher Scientific (Waltham, MA): Alexa Fluor 488-conjugated goat anti-rabbit IgG (A11034), Alexa Fluor 568- goat anti-rabbit IgG (A11036), Alexa Fluor 647- goat anti-rabbit (A21245), Alexa Fluor Plus 405- goat anti-rabbit (A48254), Alexa Fluor 568- goat anti-mouse IgG (A11031), Alexa Fluor 647- goat anti-mouse (A21235) and Alexa Fluor 647- goat anti-rat (A21247). HRP- linked secondary antibodies for immunoblotting were obtained from Jackson ImmunoResearch Laboratories (West Grove, PA): Rabbit IgG (711- 035-152), Mouse IgG (711-035-150), Rat IgG (712-035-150), and Goat IgG (705-035-003).

### Immunolabeling of brain sections

Immunoperoxidase and immunofluorescence IHC were performed according to the protocols previously described [35, 80]. Brain vibratome sections (40 μm) were blocked and incubated in primary antibody O/N (up to 3 days in some cases) at 4°C. ABC detection method was used for immunoperoxidase labeling visualized with DAB while Alexa-Fluor conjugated secondary antibodies were used for immunofluorescence. Autofluorescence was quenched with 1% Sudan black (Sigma-Aldrich; St. Louis, MO) in 70% ethanol for 20 minutes. DAB labeled sections was inspected on a Zeiss AxioSkop II equipped with a HrM digital camera (Carl Zeiss, Germany). Immunofluoescently labeled sections were examined on a Zeiss LSM880 confocal microscope.

### Confocal image collection and hue-angle based quantitative analysis for AV/LY subtypes

Confocal imaging was performed using a plan-Apochromat 20x or 40x/1.4 oil objective lens on a LSM880 laser scanning confocal microscope with the following parameters: eGFP (ex: 488, em: 490-560 with MBS 488), mRFP (ex: 561, em: 582-640 with MBS 458/561), Alexafluor 647 (ex: 633, em: 640-710 with MBS 488/561/633) and DAPI (ex: 405, em: 410-483), with the ‘best signal scanning mode’ which separates scanning tracks for each excitation and emission set to exclude crosstalk between each fluorophore signal. The resolution of 40x images was 1024 x 1024 pixels (corresponding to an area of 212.34 x 212.34 μm^2^), and the resolution of 3x digitally zoomed images was also 1024 x 1024 pixels (corresponding to an area of 70.78 x 70.78 μm^2^). Detailed settings for image collection were reported previously [35].

Hue angle-based vesicle analysis enables quantitative determination of AV/LY subtypes including AP, AL, poorly acidified AL (pa-AL) and pure LY and the method was described in detail previously [35]. Briefly, high resolution confocal images containing the LC3 (red and green) and CTSD (blue) punctate signals were analyzed with the Zen Blue Image Analysis Module from Carl Zeiss Microscopy. Threshold for each of the three-color channels (red, green, blue; RGB) was set by taking the average of intensity value from 20 neuronal perikarya. The signal was segmented into discrete puncta by using the automatic watershed function to separate clumped vesicles into individual puncta. Background signal was eliminated using the size exclusion function of Zen Blue. The R, G and B intensity values of each vesicle were calculated using the profile function of Zen Blue. The RGB ratio of each vesicle was converted into a hue angle and saturation range – which we assigned to each AV subtype and should match the desired AV color range as perceived visually (i.e., yellow for AP, blue for Ly, purple for AL and white for de-acidified AL) – by entering the values of R, G, and B for a given punctum into the formula, as follows: Hue° = IF(180/PI()*ATAN2(2*R-G-B,SQRT(3)*(G-B)) < 0,180/PI()*ATAN2(2*R-G-B,SQRT(3) *(G-B)) + 360,180/PI()*ATAN2(2*R-G-B,SQRT(3)*(G-B))). Saturation percent of the hue angle was calculated by entering the values of R, G, and B for a given punctum into the following formula = (MAX(RGB)-MIN(RGB))/SUM(MAX(RGB)+MIN(RGB))*100 (www.workwithcolor.com), provided lightness is less than 1, which is the usual case for our data. The Hue angle was converted to color using the Hue color wheel. The data was pooled and categorized in Excel spreadsheets.

### Ultrastructural analyses

Vibratome brain sections (50 μm) were treated with 1% osmium tetroxide in 100 mM sodium cacodylate buffer pH 7.4 for 30 minutes, washed in distilled water four times (10 min/wash), and then treated with 2% aqueous uranyl acetate overnight at 4°C in the dark. Samples were then washed and sequentially dehydrated with increasing concentrations of ethanol (20, 30, 50, 70, 90, and 100%) for 30 min each, followed by three additional treatments with 100% ethanol for 20 min each. Samples were then infiltrated with increasing concentrations of Spurr’s resin (25% for 1 h, 50% for 1 h, 75% for 1 h, 100% for 1 h, 100% overnight at room temperature), and then incubated overnight at 70°C in a resin mold. For TEM ultrastructural analysis, 70 nm sections were cut using a Leica Reichert Ultracut S ultramicrotome and a Diatome diamond knife, placed on to grids and then post stained with 2% uranyl acetate and lead citrate. Images were taken using a Ceta Camera on a ThermoFisher Talos L120C transmission electron microscope operating at 120kV.

For post-embedding IEM, ultrathin sections were placed on to carbon formvar 75 mesh nickel grids and etched using 4% sodium metaperidotate for 10 minutes before being washed twice in distilled water and then blocked for one hour. Grids were incubated in the primary antibodies at 4 °C overnight. Next day grids underwent seven washes in 1xPBS and were then incubated in anti-mouse or anti rabbit 10 nm gold secondary (1/50 dilution) for 1 hour. After this the grid was washed seven times in 1xPBS and twice in distilled water. The grids were then silver enhanced for 5 minutes (Nanoprobes). Grids were finally post stained with 1% uranyl acetate for 5 minutes followed by two washes in water and then stained with lead citrate for 5 minutes followed by a final two washes in distilled water. Samples were then imaged on a ThermoFisher Talos L120C operating at 120kV.

### Western blotting

Samples for WB were prepared by homogenizing brains in a tissue- homogenizing buffer (250 mM sucrose, 20 mM Tris pH 7.4, 1 mM EDTA, 1 mM EGTA) containing protease and phosphatase inhibitors as previously described. [67] Following electrophoresis, proteins were transferred onto 0.2 µm-pore nitrocellulose membranes (Whatman, Florham Park, NJ) at 100 mA for 8-12 hours depending on the target protein. The blots were blocked for 1 hour in 5% non-fat milk in TBS, rinsed in TBST (TBS + 0.1% Tween-20), then incubated with a primary antibody in 1% BSA/TBST overnight at 4°C. The membrane was washed and incubated in a HRP conjugated goat-anti rabbit or mouse secondary antibody, diluted 1:5000 in 5% milk for 1 hr at room temperature. The membrane was again washed and then incubated in a Novex ECL (Invitrogen) for 1 min. The detection of the signals was achieved through either exposure to a film or scanning by a digital gel imager (Syngene G:Box XX9) as specified in the figure legends. Densitometry was performed with Image J and the results were normalized by the immunoblot(s) of given loading control protein(s) (usually GAPDH, unless otherwise noted).

## Results

### Identification of inclusions and HTT molecular species in Q175 mice

Immunohistochemistry (IHC) experiments with the antibody mEM48, which preferentially recognizes aggregated mHTT [20], revealed age-dependent development of mHTT-positive profiles in the striatum of Q175KI mice. Brain sections from 2.5-mo-old Q175 (not shown) only exhibited faint and diffuse nuclear mHTT staining without identifiable aggresomes/inclusion bodies (IBs), while at 6 and 10 months of age, mHTT-positive IBs were readily detected progressively with age (Fig. 1A2-A3). There were no similar mHTT immunoreactive puncta in WT striatum (Fig. 1A1). These observations are consistent with previous findings in Q175 and other HD mouse models [12, 38, 48]. To further determine the locations of mHTT IBs in Q175, mEM48-immunostained sections were counterstained with cresyl violet (Fig. 1A4) to distinguish nuclear IBs (Fig. 1A4, arrowheads) from extranuclear IBs, which were localized predominantly in the neuropil (Fig. 1A4, arrows) and detected, but rarely, in the cytoplasmic portion of the perikaryon. Thus, our results demonstrate age-dependent increase in mHTT aggresomes in the Q175 model.

**Fig. 1.**
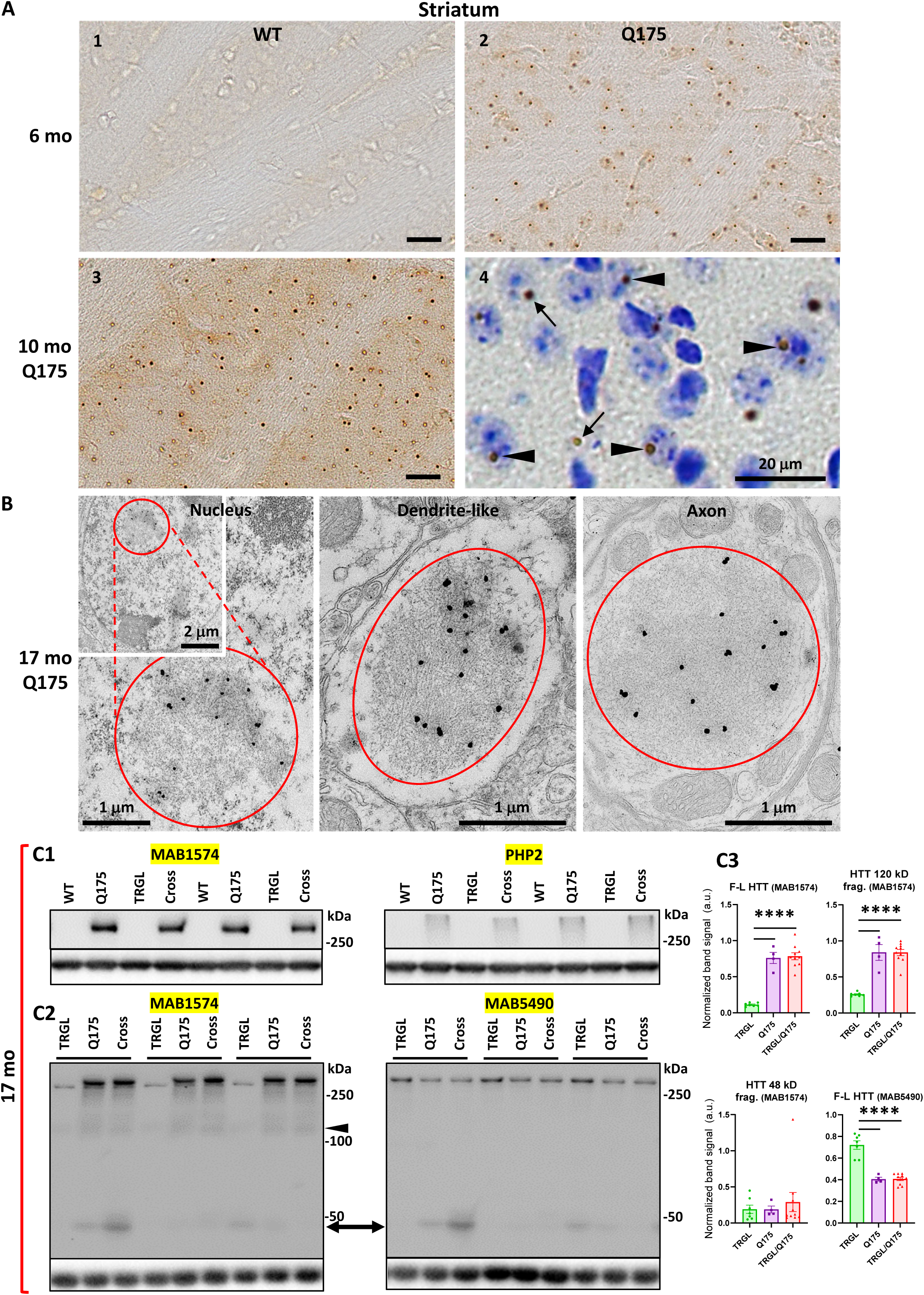
Identification of inclusions and HTT molecular species in Q175 mice. ***(A) IHC detects age-dependent increase in the number of mHTT inclusions.*** Brain sections from 6-mo- and 10-mo-old WT and Q175 mice were processed for IHC with antibody mEM48 (MAB5374) directed against mHTT (A). Images from sections without (A1-3) or with (A4) a cresyl violet counterstain are shown where the dark-brown puncta represent mHTT-positive inclusions. Arrowheads depict neuronal intranuclear inclusions (NIIs), determined with the assistance from the nuclear labeling by cresyl violet, while arrows indicate extranuclear inclusions, primarily the neuritic inclusion in the neuropil. Bars = 20 μm. n = 4 mice/genotype, 4 sections/mouse. ***(B) mHTT inclusions are detected in nucleus, dendrites and axons of Q175 brains by IEM.*** Sagittal vibratome brain sections of 17-mo-old Q175 were cut and went through EM processing. Small blocks were obtained from the striatal areas for ultrathin sectioning. Tissue containing grids were processed for immunogold labeling procedure with antibody mEM48, using 10 nm gold followed by silver enhancement. Structures showing high level of silver-enhanced gold labeling were considered as mHTT-positive. ***(C) Various forms of HTT molecules are detected with different antibodies by immunoblotting.*** Equal amounts of proteins from hemibrain homogenates of 17-mo-old WT, TRGL, Q175 and TRGL/Q175 (labeled as “Cross”) were subjected to SDS-PAGE and processed for WB with different antibodies directed against HTT/mHTT, including MAB1574 (C1, C2), mAb PHP2 (C1) and MAB5490 (C2). Images were collected by a digital gel imager (Syngene G:Box XX9). The arrowhead and arrow (C2) depict a 120 kDa and a 48 kDa fragment, respectively. (C3) Densitometry was performed with Image J for the blots shown in (C2) and the results were normalized by the immunoblot(s) of given loading control protein(s) (e.g., GAPDH). Values are the Mean +/- SEM for each group (n = 7 TRGL, 4 Q175 and 10 TRGL/Q175). Significant differences among the groups were analyzed by One-Way ANOVA followed by Sidak’s multiple comparisons test. * P < 0.05, ** P < 0.01.

To reveal the ultrastructural locations and features of the mHTT aggregates, immunogold EM (IEM) with antibody mEM48 was performed. EM images (Fig. 1B) demonstrated that the IEM with this antibody was highly specific in detecting mHTT IBs which were localized in the neuronal nuclei, dendrites and axons. Ultrastructurally, most IBs were cotton-ball shaped and composed of fine fibrous or granular elements, somewhat similar to the unbundled short fibrils/protofibrils found *in vitro* with recombinant mHTT protein fragments [32, 33, 44], and similar to the structure of NII type inclusions we found in the human HD brain [9]. Notably, however, human neuritic IBs often displayed a more heterogeneous composition, such as fine fibrils mixed with AVs or bundles composed of microtubule-like filaments, which were not found in the mouse neuritic IBs. Thus, the result suggests a homogenous aggresome pathology in the mouse model, possible reflecting a more rapid formation rate than in human.

Previous immunoblotting studies have observed fragmentation of mHTT molecules in the human brain [30, 49], including our own study which detects mHTT fragments of 45-48 kDa, which predominantly exist in HD striatum [9]. To assess HTT molecular species in the Q175 mouse brain, we employed multiple antibodies that preferentially detected mHTT over wild type HTT, including MAB1574 (Clone 1C2) (Fig. 1C1, C2), its epitope containing a 38-glns stretch [72], and mAb PHP2 (Fig. 1C1), reacting with the peptide sequence QAQPLLPQP within the proline rich domain of HTT [32]. Both detected full-length mHTT and a ∼120 kDa fragment in the Q175 model (Figs. 1C1; 1C2 left and 1C3 top two graphs). By contrast, MAB5490 (Fig. 1C2 right), reacting with aa115-129 of HTT (C-terminal to the region of polyQ stretch-containing exon 1), detected both wild type and mutant forms of HTT (Fig. 1C3, bottom right graph). A ∼48 kDa HTT fragment may correspond to the 45-48 kDa fragment seen in human brain [9], and was detected by both MAB1574 and MAB5490 antibodies in some samples (Fig. 1C2) but its levels in the Q175 models and the control TRGL were not statistically significantly different (Fig. 1C3, bottom left graph). Thus, the result suggests that HTT fragmentation is not obvious in the Q175 brains, unlike the far more prevalent occurrence of this phenomenon in human HD striatum.

### mHTT colocalizes with p62, Ub and CTSD in Q175 striatum

Our earlier study in human HD brains [9] found a high degree of colocalization of mHTT/Ub or Ub/p62 colocalized signals in IBs, suggesting a relationship between mHTT and the autophagy machinery since p62 and Ub are adaptor proteins mediating autophagic cargo sequestration. Similarly, in Q175 mice, mHTT/Ub or Ub/p62 signals were highly colocalized in IBs, particularly in NIIs, in striatal neurons of Q175 mice (Fig. S1). Additional triple IF labeling experiments (mHTT/Ub/p62) with brain sections from 6- and 10-mo-old Q175 mice identified IBs positive for mHTT, p62 and Ub within or outside the nuclei (Fig. 2A, arrows and arrowheads, respectively). The 10-mo-old Q175 mice exhibited more and larger mHTT-positive IBs than 6-mo mutants (Fig. 2A, 1^st^ column), consistent with Fig. 1A and the literature [12, 16]. Thus, our data demonstrate a close spatial relationship among mHTT and autophagy receptor protein p62 and Ub in the Q175 model, which is consistent with our observations in human HD brain [9].

**Fig. 2.**
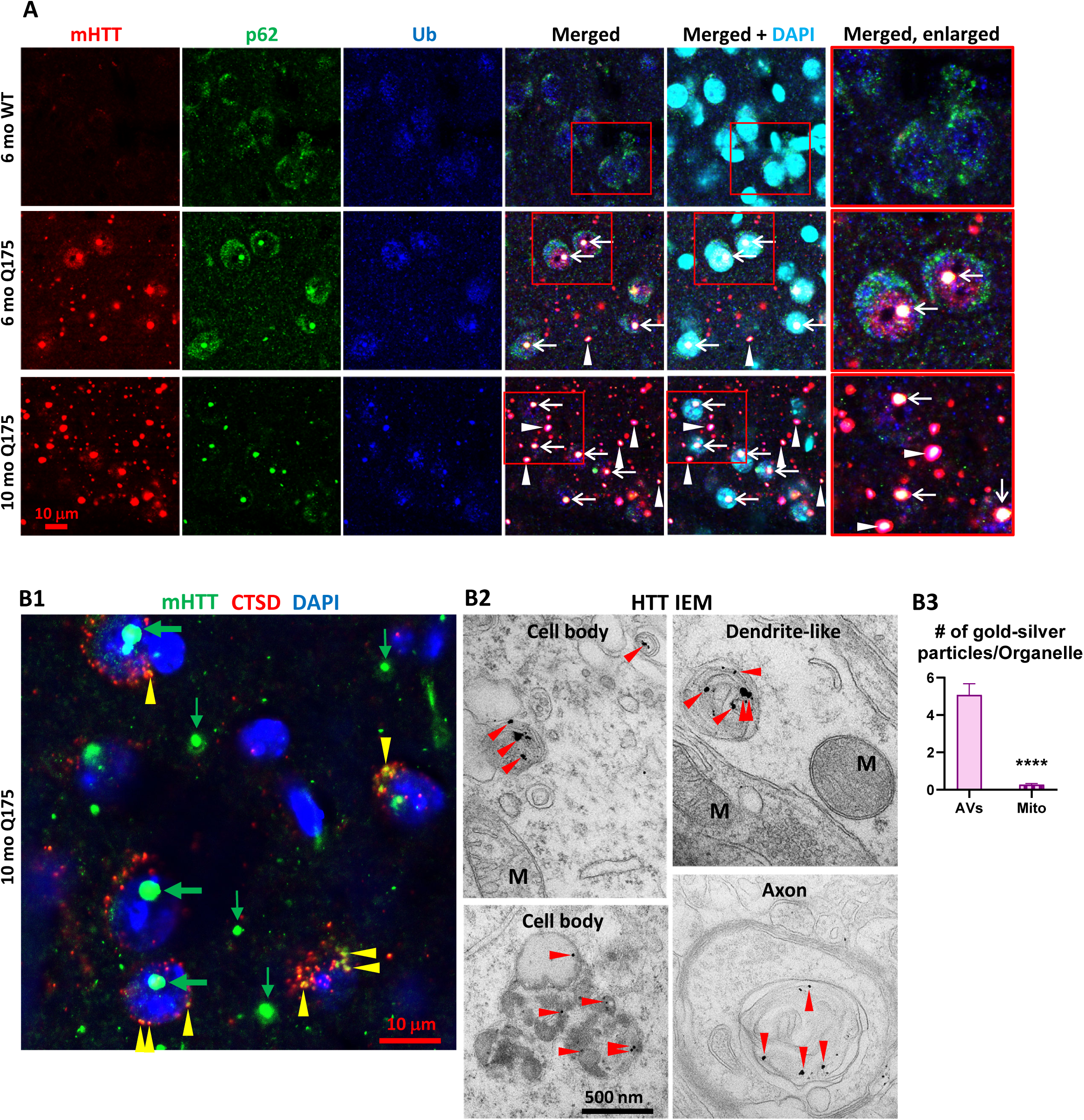
Colocalization of mHTT with p62, Ub and CTSD. ***(A) Triple labeling detects colocalization of mHTT with both autophagy adaptor proteins p62 and Ub.*** Brain sections from 6-mo- and 10-mo-old mice were immunostained with antibodies to HTT (antibody MW8; red), p62 (green) or pan-Ub (blue), followed by an additional DAPI (cyan) labeling, and confocal images from the striatum are shown. Boxed areas are enlarged and shown in the last column. Arrows and arrowheads depict the white areas representing triply labeled inclusions containing the signals of the three proteins, where arrows are for NIIs, determined with the assistance from the nuclear DAPI labeling, while the arrowheads are for extranuclear inclusions. Bar = 10 μm. n = 4 mice/genotype, 4 sections/mouse. ***(B) mHTT is detected in vesicles of the ALP.*** (B1) Brain sections from 10-mo-old Q175 were double-immunolabeled with antibodies to HTT (antibody MW8; green) and CTSD (red), followed by an additional DAPI (blue) labeling, and a 3-color-merged confocal image from the striatum is shown. Large and small green arrows depict HTT nuclear and extranuclear inclusions, respectively, while yellow arrowheads depict yellow puncta showing HTT and CTSD signal colocalization. Bar = 10 μm. n = 4 mice/genotype, 4 sections/mouse. (B2) IEM with anti-HTT antibody (MAB1574) specifically detects HTT signal, represented by the silver-enhanced gold particles (red arrowheads), in AV/LY in cell bodies, dendrites and axons. (B3) To demonstrate the labeling specificity of the HTT antibody in this IEM study, the number of silver-enhanced gold particles in AV/LY existing in neuronal cell bodies and neurites was counted from 69 EM images from two 10-mo-old Q175 mice against the number of silver-enhanced gold particles in mitochondria on the same images, and the result is shown in the bar graph. Statistical significances between the two groups were analyzed by unpaired, two-tailed t-test. **** P < 0.0001.

Double IF labeling of mHTT with a LY marker cathepsin D (CTSD) (Fig. 2B1) detected small punctate mHTT signal in CTSD-positive vesicles, suggesting a pool of mHTT within the ALP. Consistent with this LM finding, IEM with antibody MAB1574 directed against mHTT clearly demonstrated that the AVs in either cell bodies, dendrites and axons were labeled with concentrated silver-enhanced gold particles (Fig. 2B2), and the specificity of the IEM labeling was very high, as reflected by a much higher number of silver-enhanced gold particles associated with AVs versus the minimal number of silver-enhanced gold particles with mitochondria that represent the background labeling (Fig. 2B3). Together, the LM and IEM findings suggest that mHTT molecules exist within ALP vesicles in the absence of definable aggresome/IB structures, similar to the IEM finding from a recent report [82].

### Mild late-onset alterations in the ALP revealed in 17-mo-old Q175 mice

We crossed TRGL mice [35] with Q175 to generate TRGL/Q175 and assessed AV/LY subtypes in striatal neurons with a hue angle-based analysis method [35, 37]. Our pilot studies in young TRGL/Q175 mice (2.5 to 6-mo of age) hardly detected autophagy alterations, compared to the control TRGL mice (not shown). Previous studies have reported that autophagy impairment was hardly detected in 6-mo Q175/GFP-LC3 mice [78], but alterations in a limited number of autophagy markers (e.g., protein levels of p62, LC3, p-Beclin-1) were found in brain homogenates of 12-15-mo Q175 mice [1, 21, 78]. Therefore, to further validate the status of the ALP and to discover potential additional autophagy alterations, we expanded our study to old mice at 17-mo of age, where we found the following ALP alterations.

***(1) Alterations in AV/LY subtypes.*** Confocal images from 17-mo-old TRGL/Q175 and control TRGL brain sections, triple-fluorescent labeled via an additional immunostaining with an anti-CTSD antibody and far-red emitting fluorophore, showed enhanced mRFP- and eGFP-LC3 signals in the striatal neurons of TRGL/Q175 compared to TRGL (Fig. 3A1). Hue angle-based analysis confirmed the above observations by revealing significant increases in the numbers of AL and poorly-acidified AL (pa-AL, as explained in Lee et al., 2019; 2022). It also detected a reduction in the number of LY (Fig. 3A2). Thus, these results indicate that the tfLC3 reporter can reveal alterations in the proportions of AV/LY subtypes in Q175, consistent with delayed and/or deficient acidification of AL causing deficits in the reformation of LY to replenish the LY pool.
***(2) Mild alterations in the levels of autophagy marker proteins detected by immunoblotting.*** To further assess autophagy phenotypes in Q175 models, we conducted western blotting (WB) using hemibrain homogenates from 17-mo-old mice for protein markers of individual steps in the ALP (i.e., autophagy induction, membrane nucleation/AP formation, autophagy adaptor proteins and AL formation/substrate degradation) (Fig. S2). At the stage of autophagy induction signaling, we did not see alterations in the levels of mTOR and p-mTOR forms including its auto-phospho-form at S2481, but we did observe a decreased level of a mTOR substrate, p-p70S6K (T389), in TRGL/Q175 compared to the control TRGL (Fig. S2A), implying reduced mTOR activity, similar to the finding from a previous study [59]. Another mTOR substrate, p-ULK1 (S757), did not exhibit changes in TRGL/Q175. However, there was an increase in the level of total ULK1, resulting in a reduced ratio of p-ULK1 (S757)/total ULK1 in TRGL/Q175 compared to TRGL (Fig. S2A). The data collectively imply a down-regulated mTOR activity, as expected in response to accumulated aggregate-prone protein in Q175 mice, or as a result of sequestration of mTOR by mHTT IBs as suggested previously [59].

**Fig. 3.**
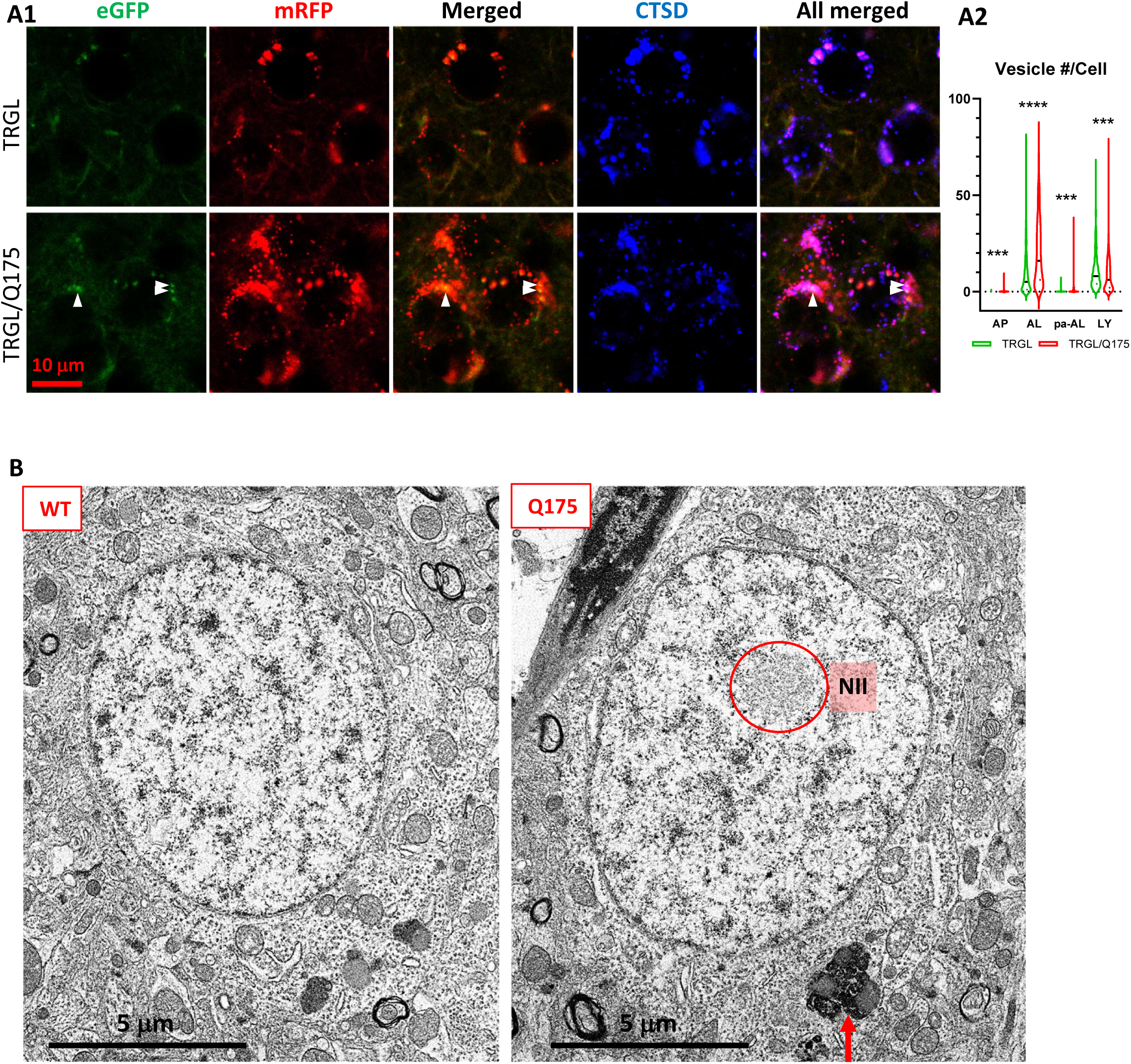
Mild late-onset alterations in the ALP in the striatum of 17-mo-old Q175. ***(A) Quantitation of AV/LY subtypes of striatal neurons detect increases in AL, pa-AL and a decrease in LY in 17-mo-old TRGL/Q175 vs. TRGL.*** (A1) Brain sections from TRGL and TRGL/Q175 (4 sections/mouse, 10 mice/genotype) were immunostained with an anti-CTSD antibody. Confocal images from the cranial-dorsal portion of the striatum (3 images at 120x/section) were collected and representative images for each eGFP-LC3 (green), mRFP- LC3 (red) and CTSD (blue) are shown. Arrowheads depict pa-AL. (A2) Hue angle-based analysis was performed for AV/LY subtype determination using the methods described in Lee et al., 2019 (see the Materials and Methods). Data are presented as Vesicle #/Neuron (TRGL: n = 713 neurons; TRGL/Q175: n = 601 neurons). Statistical significances between the two groups for each vesicle type were analyzed by unpaired t-test. Two-tailed P value: *** P<0.001, **** P<0.0001. ***(B) EM detects larger AL/lipofuscin granules in the Q175 striatum.*** Sagittal vibratome brain sections from 17-mo-old mice were cut and went through EM processing. Small blocks were obtained from the striatal area for ultrathin sectioning, followed by EM examinations of the grids. The red circle depicts NII, and the arrow depicts larger sized (> 1µm) lipofuscin granules, which were counted on randomly collected images of neurons from striatum of WT (420 neurons from n = 6 mice) or Q175 (586 neurons from n = 9 mice).

Except for these alterations, we did not detect statistically significant differences between TRGL/Q175 and the control TRGL in all other tested marker proteins in the downstream phases of the ALP (Fig. S2B-D). Of special interest is the ATG14-containing VPS34/Beclin-1 complex implicated in HD pathogenesis [54, 78] for which we did not detect statistically significant alterations in the levels of Beclin-1, VPS34 and ATG14 and their corresponding phosphor-forms. Notably, even if some TRGL/Q175 clearly exhibited diminished signals for p-ATG14 (S29) (Fig. S2B), no statistical significance could be established due to large variations among samples (See Discussion). Another notable point is that we did not find alterations in levels of full-length proteins or fragments [25, 51, 64, 73] of autophagy adaptor protein such as p62, TRAF6 (Fig. S2C), in contrast to our immunoblot analysis of human HD brains [9]. Together, our data suggest that, in general, autophagy alterations at the protein level, as can be detected by immunoblotting, are mild in TRGL/Q175 even at 17-mo-old (See Discussion).

***(3) Increased numbers of larger lipofuscin granules.*** To assess possible ultrastructural pathologies, we conducted EM study for Q175 vs WT at 17-mo of age. As predicted, NIIs were not observed in WT neurons (Fig. 3B, left panel) but were detected (Fig. 3B, right panel, red circle) in ∼10% striatal neurons of Q175 mice. Different from the substantial accumulation of AVs including lipofuscin granules in late stage human HD brain (e.g., HD4) [9], we did not observe gross accumulation of AVs in Q175 at this old age, except that quantitative analysis of EM images revealed that larger sized (>1 µm) lipofuscin granules (Fig. 3B, right panel, arrow) modestly increased in number in Q175 compared to WT [WT: 113 lipofuscin/420 neurons (n = 6 mice), i.e., 27 lipofuscin/100 neurons; Q175: 228 lipofuscin/586 neurons (n = 9 mice), i.e., 39 lipofuscin/100 neurons; Unpaired t-test, Two-tailed P value < 0.01; Bar graph not shown]. Thus, these EM findings suggest that ultrastructural autophagy alterations in the ALP were mild in Q175 even at an older age.

### DARPP-32-IR decreases in striatal neurons of 17-mo-old TRGL/Q175 in the absence neuronal loss

To investigate potential neurodegeneration in our model, brain sections from 17-mo-old mice were double-labeled with the medium spiny neuron marker DARPP-32 (dopamine- and cAMP-regulated phosphoprotein, 32 kDa)[53] and a general neuronal marker NeuN. We found that the intensity of DARPP-32-IR significantly diminished in TRGL/Q175 compared to TRGL (Fig. 4A, B). However, there were no alterations in NeuN-IR, including Area Covered and the Number of NeuN-positive neurons (Fig. 4A, B), and there were clear examples of neurons showing strong NeuN signal but faint or no DARPP-32 signal (Fig. 4A, arrows). Such a finding is consistent with other studies in Q175 and Q140 mice where neuronal loss was not found even though a reduction in DARPP-32 was seen in the same study, and that neuronal loss only occurred quite late, e.g., around 2 years of age [16, 22, 55, 62]. Thus, our data support the notion that changes in DARPP- 32 may indicate alterations in its protein level and a loss of phenotype pattern, which is a potentially common prelude to neuronal loss in other neuronal cell types, e.g., basal forebrain cholinergic neurons [27].

**Fig. 4.**
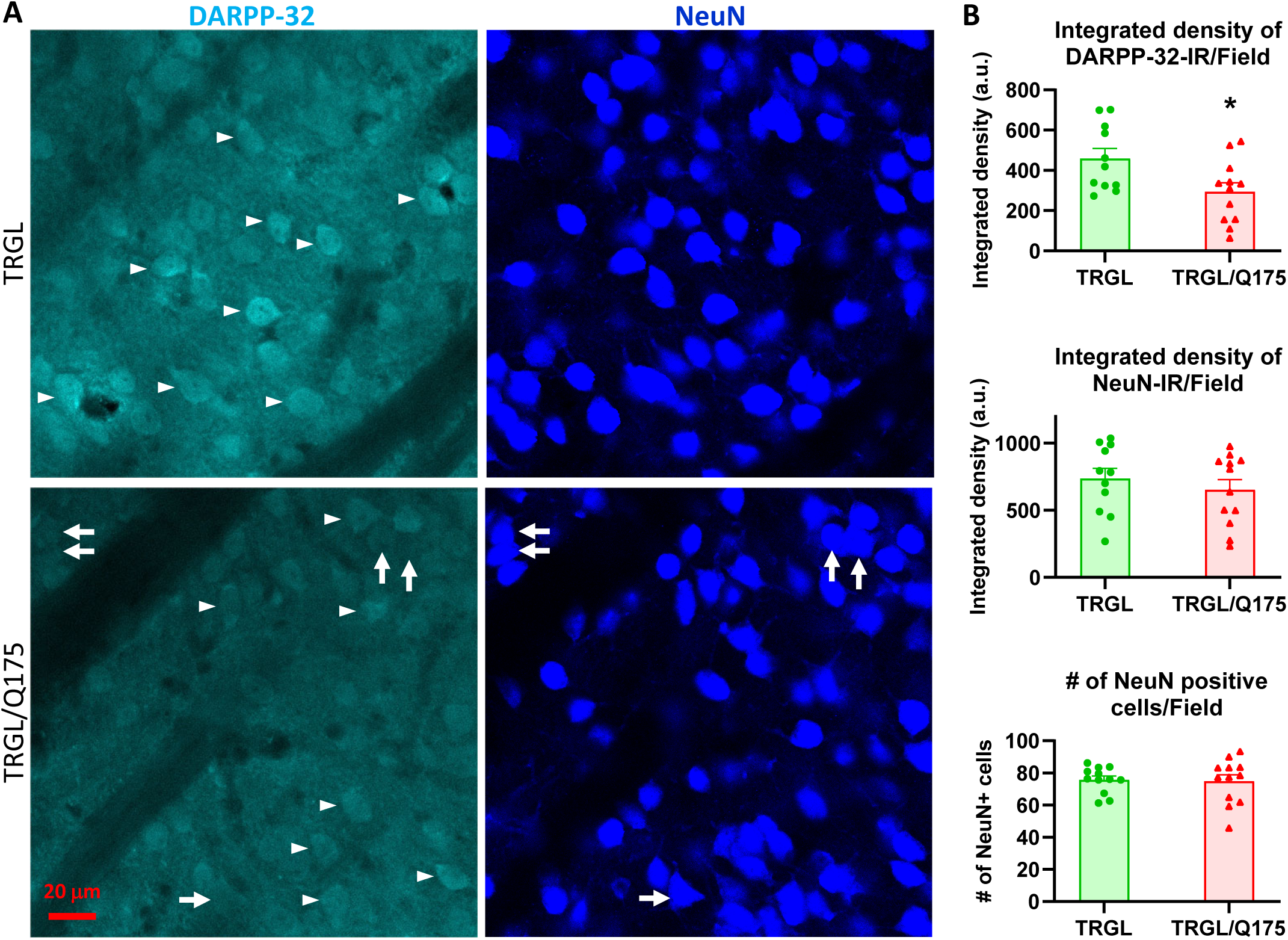
Decrease of DARPP-32-IR in striatal neurons of 17-mo-old TRGL/Q175 in the absence of a NeuN signal reduction. Brain sections from TRGL and TRGL/Q175 (n = 10 mice/genotype, 4 sections/mouse) were double-immunostained with anti-DARPP-32 and -NeuN antibodies. Confocal images from the cranial-dorsal portion of the striatum (3 images at 120x/section) were collected and representative images are shown (A). Arrows depict examples of neurons showing strong NeuN signal while minimal DARPP-32-IR. (B) Images were quantified by Image J for Integrated Density of DARPP-32- (top) or NeuN-IR/Image field (middle), and for # of NeuN positive cells/Image field (bottom). Statistical significances between the two groups were analyzed by unpaired, two-tailed t-test. * P < 0.05.

### Oral mTORi INK engages mTOR target and induces downstream responses in the ALP in 7-mo-old TRGL/Q175

As a prelude to investigate effects of pharmacological autophagy stimulation with a mTORi, INK, on TRGL/Q175 mice, we first performed a safety, pharmacokinetic and pharmacodynamic evaluation of INK in 6-mo-old WT mice, administered via oral gavage. The WB results (Fig. S3A) indicated that all doses tested, from 2.5 mg/kg (mpk) to 10-mpk, exhibited target engagement as revealed by the reductions in the protein levels of mTOR targets, i.e., p-ULK1 (S757, phosphorylated by mTOR), p-p70S6K (not shown) or phosphorylated S6 ribosomal protein (p-S6, at S240/244), implying high BBB permeability of this compound and target engagement. Consistently, measurements for the levels of INK in cerebellar homogenates of WT mice treated with 1, 3 and 10-mpk INK revealed dose-dependent brain levels of INK (Fig. S3B).

We then decided an oral gavage treatment regimen for INK as 4-mpk, daily, for 3 weeks, to be administered in 6-mo-old TRGL/Q175. The brains from the mice (7-mo of age after the 3-week treatment duration) were then analyzed by multiple experimental approaches. By immunoblotting, target engagement was verified, as indicated by decreased levels of p-mTOR (S2481, autophosphorylation site), p-ULK1 (S757) and p-S6 (S240) in brain homogenates from INK-treated TRGL/Q175 compared to Veh-treated TRGL/Q175 (Fig. 5A, B). Additionally, there were also INK induced changes in marker proteins located downstream of autophagy induction, including, for example, increased levels of p-ATG14 in INK-treated samples (Fig. 5A, B). It is interesting that although INK did not induce any alterations in the transgene product tfLC3-I or - II, there was a trend of decreased endogenous LC3-I, leading to a statistically significant increase in the ratio of LC3-II/I, and such a result in LC3-I reduction was reproduced in a repeated experiment (Fig. 5A, B). Thus, the pilot study in WT mice and the actual study in TRGL/Q175 mice together established mTORi INK’s BBB permeability, target engagement and ability to modify molecular events in the ALP.

**Fig. 5.**
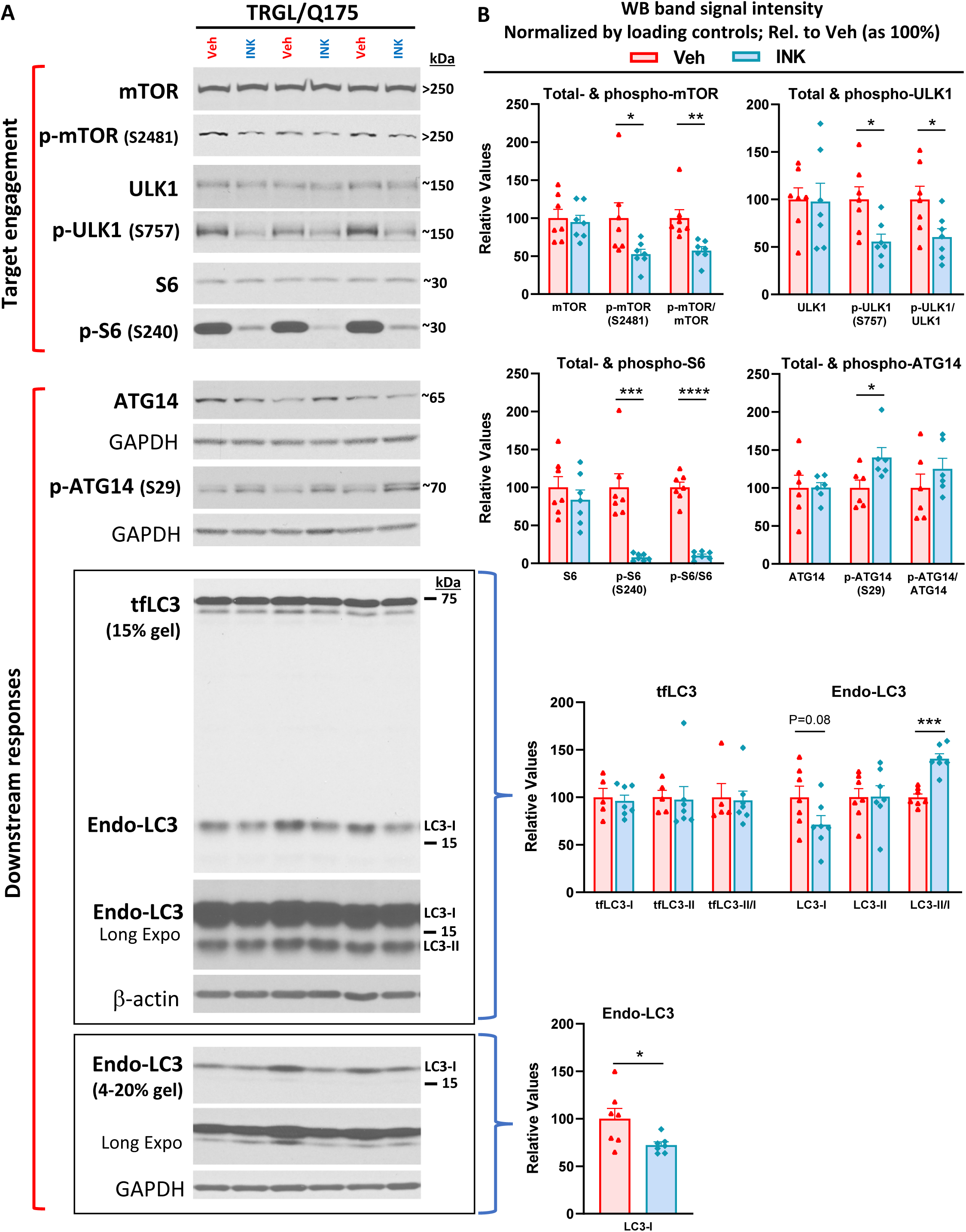
mTORi INK exhibits target engagement and induces downstream responses in the ALP in 7-mo-old TRGL/Q175. Equal amounts of proteins from brain homogenates of 7-mo-old TRGL/Q175 mice untreated (labeled as “Veh”) or mTORi INK treated [4 mg/kg (4-mpk), daily, 3 weeks; labeled as “INK”] were subjected to SDS-PAGE and processed for WB with antibodies directed against several marker proteins in the autophagy pathway, representing target engagement of INK or downstream responses. Representative blots are shown on the left (A) while quantitative results of the blots are shown on the right (B). The bottom LC3 blots represent a repeated immunoblotting experiment. Values are the Mean +/- SEM for each group (n = 6-7 mice per condition). Significant differences between the two groups were analyzed by unpaired, two-tailed t-test. * P<0.05, ** P<0.01, *** P<0.001, **** P<0.0001. tfLC3 = mRFP-eGFP-LC3; Endo-LC3 = endogenous LC3.

### mTOR inhibition alters AV/LY subtypes

We took advantage of the tfLC3 construct reporting *in vivo* autophagy flux [35] to assess INK effects on AV/LY subtypes in TRGL/Q175. Confocal images from INK- or Veh-treated TRGL/Q175 and Veh-treated TRGL brain sections of 7-mo-old mice, immuno-stained with anti-CTSD antibody and detected with a third fluorophore, were collected (Fig. 6A) and processed for hue-angle based quantitative analysis using the protocol described previously [35]. The analysis in striatal neurons (Fig. 6B) revealed reversal of the two main ALP alterations. First, the pre-existing abnormally lowered LY number in TRGL/Q175-Veh (Fig. 6B, LY group, red vs green), was reversed by INK treatment (Fig. 6B, LY group, red vs blue). Second, INK reduced the number of AL in TRGL/Q175-Veh compared to TRGL-Veh (Fig. 6B, AL group, red vs blue), implying improved AL clearance and likely providing the basis for restoration of normal LY number. Together, the data suggest that autophagy phenotype in 7-mo Q175 is measurable but milder than that in the 17-mo-old mice shown in Fig. 3A and that INK has the potential to modulate the autophagy pathway and restore normal function at this mild disease stage.

**Fig. 6.**
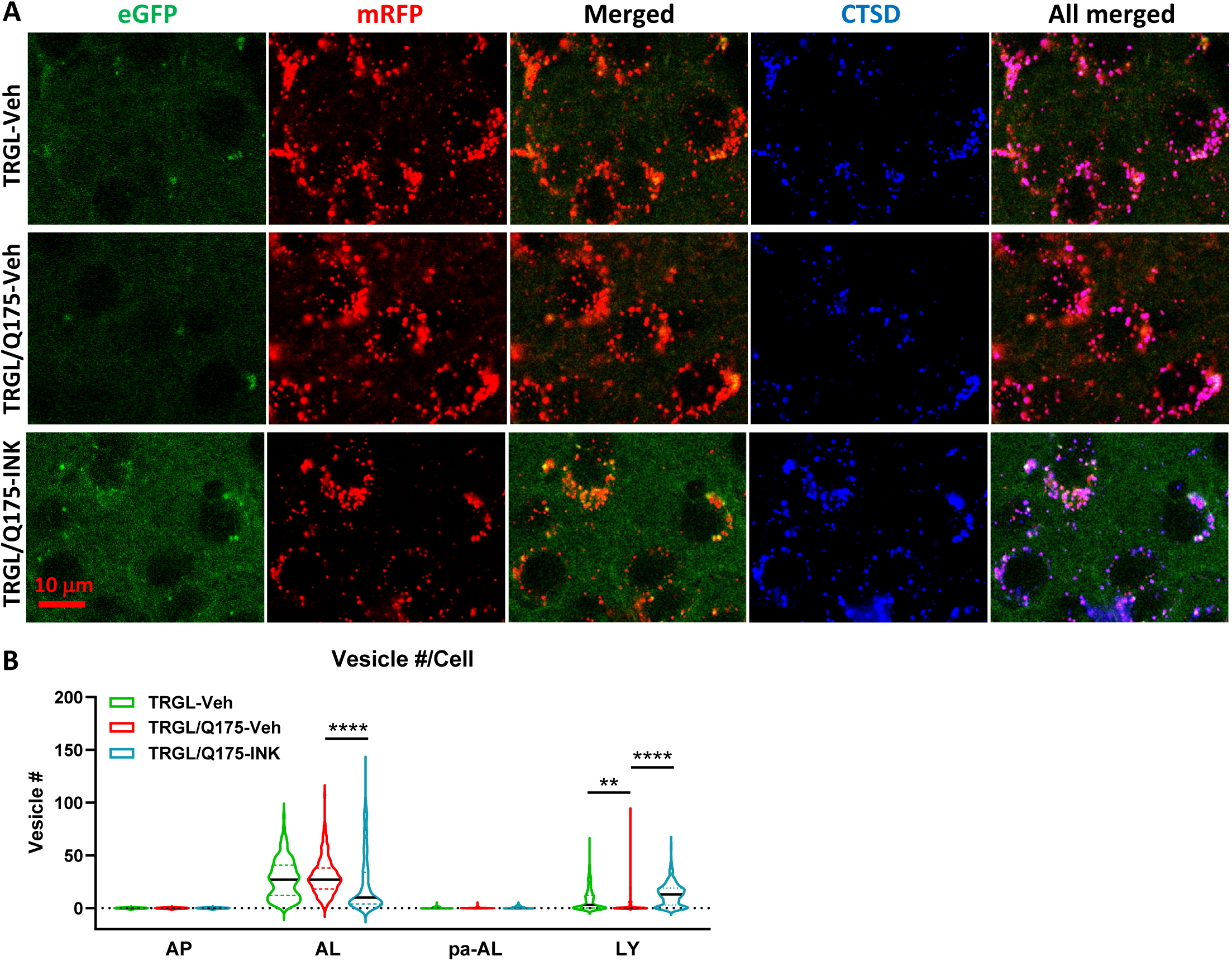
mTORi INK reverses the mild alteration of AV/LY subtypes in the striatum of 7-mo-old TRGL/Q175. (A) Brain sections from untreated or INK (4-mpk, 3w)-treated TRGL/Q175 vs. TRGL (4 sections/mouse, 5-6 mice per condition) were immunostained with an anti-CTSD antibody. Confocal images from the cranial-dorsal portion of the striatum (3 images at 120x/section) were collected and representative images for each eGFP-LC3 (green), mRFP-LC3 (red) and CTSD (blue) are shown. (B) Hue angle-based analysis was performed for AV/LY subtype determination using the methods described in Lee et al., 2019. Data are presented as Vesicle #/Neuron (TRGL-Veh: n = 260 neurons; TRGL/Q175-Veh: n = 218 neurons; TRGL/Q175-INK: n = 287 neurons). Statistical significances among the groups were analyzed by One-Way ANOVAfollowed by Sidak’s multiple comparisons test. ** P<0.01, **** P<0.0001.

### mTOR inhibition mitigates mHTT-related aggresome pathology

Importantly, we found that HD-like pathology such as mHTT IBs and associated protein aggregation of adaptor proteins, p62 and Ub, was ameliorated by INK treatment. Confocal images demonstrated that mHTT positive particles/inclusions were substantially reduced in INK-treated 7-mo-old TRGL/Q175, compared to Veh-treated TRGL/Q175 mice (Fig. 7A). Moreover, p62-IR or Ub-IR also changed in a similar trend (Fig. 7A), consistent with the high degree of colocalization of mHTT/p62/Ub signals shown in Fig. 2A. The corresponding quantitative analysis, shown as Area Covered by either mHTT-, p62- or Ub-IR on a per cell basis, confirmed statistically significant decreases in the Total Area Covered by mHTT- and p62-signals (Fig. 7B1). With the assistance of the endogenous tfLC3 signal (particularly the mRFP signal) in the TRGL to identify the association status of the mHTT-, p62- or Ub-IR with AVs, the images were additionally quantified to generate the separated “Area Covered by the AV-associated Form” (Fig. 7B2) and “Area Covered by the AV-unassociated Form” (Fig. 7B3) of mHTT-, p62- or Ub-signals. The results collectively suggest decreased mHTT-, p62- or Ub-IR in treated compared to untreated TRGL/Q175, implying that INK treatment was able to promote the clearance of mHTT and related receptor proteins p62 and/or Ub.

**Fig. 7.**
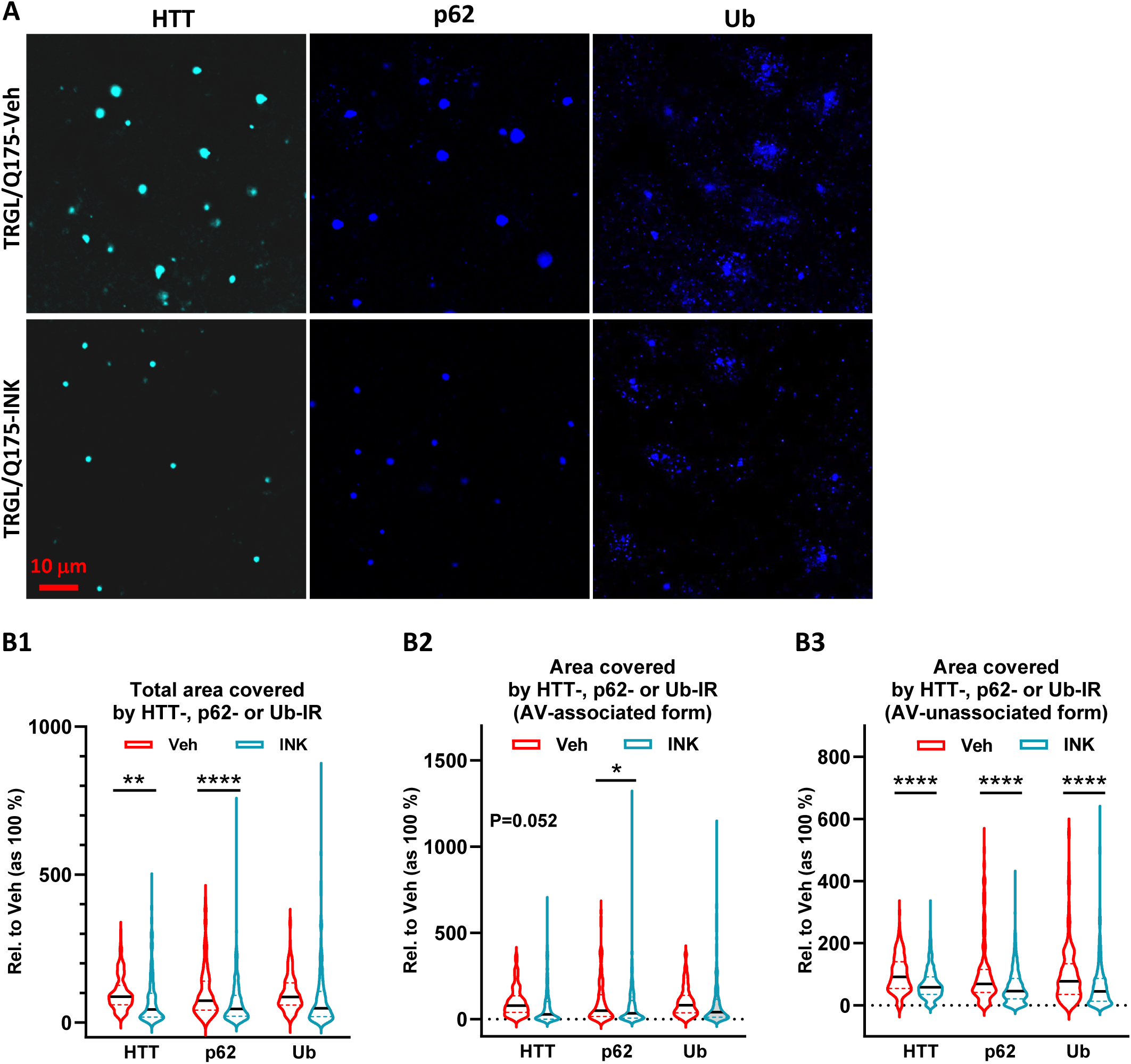
mTORi INK reduces HTT-, Ub- or p62-IR-covered areas parallelly in the striatum of 7-mo-old TRGL/Q175. (A) Brain sections from INK (4-mpk, 3w)-treated or untreated TRGL/Q175 (4 sections/mouse, 6-7 mice/condition,) were immunostained with anti-HTT (MW8), -p62 or -pan-Ub antibodies. Confocal images from the cranial-dorsal portion of the striatum (3 images at 120x/section) were collected. Shown are single-channel images (i.e., without showing the eGFP and mRFP signals). (B1-B3) Areas covered by either the HTT-, p62- or Ub-IR on a per cell basis are quantified (for HTT-IR, TRGL/Q175-Veh: n = 196 neurons, TRGL/Q175-INK: n = 385 neurons; for p62-IR, TRGL/Q175-Veh: n = 311 neurons, TRGL/Q175-INK: n = 378 neurons; for Ub-IR, TRGL/Q175-Veh: n = 244 neurons, TRGL/Q175-INK: n = 347 neurons) and grouped as “Total Area” (B1), “AV-associated Form” (i.e., the IR which was associated with tfLC3 signals representing AVs)(B2) or “AV-unassociated Form” (i.e., the IR which was not associated with the tfLC3 signals)(B3). Statistical significances between the two groups were analyzed by unpaired t-test. Two-tailed P value: * P < 0.05, **** P < 0.0001.

### mTOR inhibition does not reverse preexisting DARPP-32-IR reduction in TRGL/Q175

Finally, we evaluated neurodegenerative changes by employing DARPP-32 IHC on brain sections of Veh-treated or INK-treated TRGL/Q175. Similar to what was found in 17-mo-old mice (Fig. 4), the group of TRGL/Q175-Veh, even at this young age (7-mo), already exhibited a diminished DARPP-32-IR signal compared to that in TRGL-Veh (Fig. 8A, B). However, no differences between the untreated and INK-treated TRGL/Q175 groups were established by quantitative analysis (Fig. 8A, B). Thus, the result suggests a lack of effectiveness of INK in reversing this pre-existing phenotype, implying that earlier intervention may be necessary [43].

**Fig. 8.**
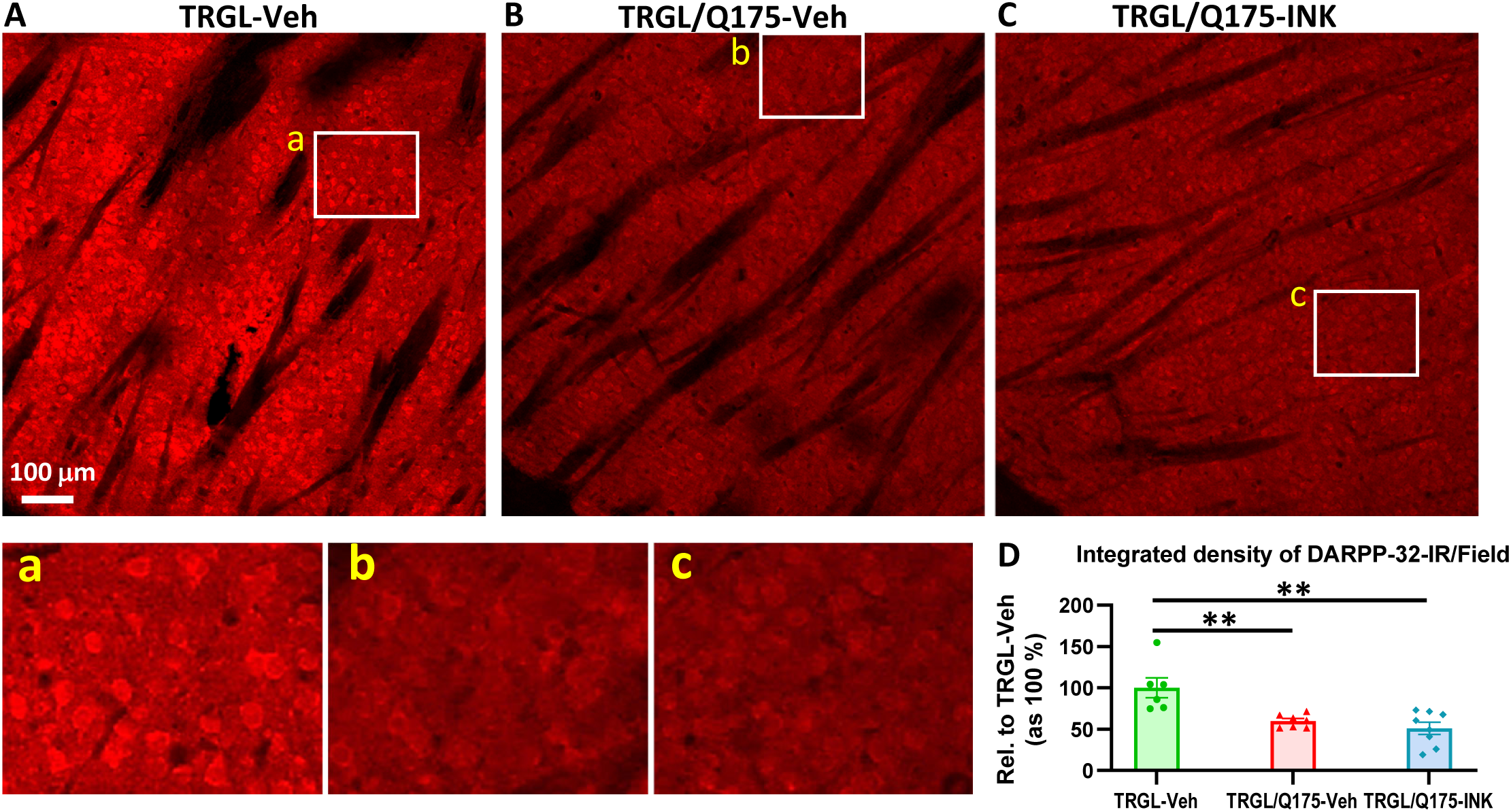
INK treatment does not reverse the reduction of DARPP-32-IR in the striatum of 7-mo-old TRGL/Q175. (A) Sagittal brain sections containing the striatum area were immunostained with an anti-DARPP-32 antibody, and 10x images taken from the cranial-dorsal portion of the striatum are shown (1^st^ row). Boxed areas on the 1st row are enlarged and shown on the 2nd row for easy viewing of the immunostaining patterns for each condition. (B) For quantitation purpose, each whole 10x image (excluding the areas covered by the fiber bundles which usually exhibited minimal background staining – achieved by threshold setting) was quantified by Image J (1 image/section, 4 sections/mouse) and the results are expressed as the Integrated Density of DARPP-32-IR. n = 6-8 mice/condition. Statistical significances among the groups were analyzed by One-Way ANOVA followed by Sidak’s multiple comparisons test. ** P < 0.01 compared to TRGL-Veh.

## Discussion

Previous studies in cell and animal models have documented roles of HTT and mHTT in autophagy in relation to HD [for reviews, see 15, 31, 45]. WT HTT participates in normal autophagy by releasing ULK1 via mTOR inhibition and serving as a scaffold to facilitate cargo sequestration by enhancing p62 interaction with LC3 and ubiquitinated cargos [52, 63]. mHTT in HD may activate autophagy by sequestering and thus inhibiting mTOR [59]. Most mHTT effects in cellular and mouse HD models appear to be inhibitory for autophagy induction steps, from initiation signaling and phagophore nucleation to cargo recognition/AP formation [3, 46, 58, 63, 78]. In addition, mHTT may disturb endolysosomal homeostasis by reducing exocytosis and promoting AL accumulation [82]. In the current study using Q175, we have found that mHTT is an ALP substrate and that the ALP is relatively intact, with some late-onset minor alterations, such as insufficient mTOR activity reflected by lowered phospho-p70S6K level and ratio of phospho- ULK1 (S757)/total ULK1, lysosome depletion and AL/pa-AL elevation. This mild late-onset nature of ALP pathogenesis allowed us to modulate ALP with a mTOR inhibitor in 6-mo-old TRGL/Q175 which resulted in positive effects on rescuing Q175 phenotypes including normalization of AL/LY subtypes and reduction in aggresome pathology.

### ALP impairments in Q175 are mild and late-onset

In the current study, we assessed ALP alterations in the Q175 model by analyzing protein markers in all phases of the ALP including autophagy induction signaling, membrane nucleation/AP formation, autophagy adaptor proteins and AL formation/substrate degradation. By immunoblot analysis, alterations in the tested marker proteins even at 17 months of age in TRGL/Q175 mice were limited to a reduction in p-P70S6K (T389), similar to that reported previously [59], and a reduced ratio of p-ULK1 (S757)/total ULK1 as a result of increased total ULK1. While these two alterations together would suggest a lower mTOR activity and autophagy activation, consequential alterations of downstream markers, particularly proteins in the ATG14- containing Beclin-1/VPS34 complex, were not detected. Although 4 out of the 10 tested TRGL/Q175 mice exhibited very low levels of p-ATG14, implying a deficit in this complex as previously reported [78], variation among animals was too great to draw statistical conclusions. In this regard, mice used in the current study were heterozygous, which results in less autophagy alteration than that reported in homozygous mice using certain autophagy markers [1]. It may also be noted in our experience that immunoblot analysis of brain tissue averages contributions from varied cell types that may respond to disease states in opposite directions and mask changes in a given cell population. Thus, we also applied a neuron-specific analysis using our neuron-specific tfLC3 reporter and hue-angle based image quantification approach, which yielded more sensitive detection and findings.

In brain sections from 17-mo-old TRGL/Q175 mice, we observed increased numbers of AL including the population of pa-AL and reduced numbers of LY. Such changes in AL/pa-AL were not readily observed in TRGL/Q175 mice at 2 months (not shown) or 7 months (e.g., Fig. 6B, which only exhibited a reduction in LY number) of age, indicating that even if these alterations signify an existence of impairments in the late phase of the ALP, they do not occur until a later age (e.g., 17 mo). Importantly, this pattern of late-onset ALP pathology progression, e.g., AL/pa-AL increases, is consistent with our findings from HD human brains [9] where enlarged CTSD-positive AL accumulate and cluster in affected neuronal populations at the late disease stages (HD3 and HD4). These findings suggest that the overall autophagy function in heterozygous Q175 is largely maintained until the relatively late disease stage and provide a rationale to stimulate autophagy as a therapeutic intervention when the autophagy machinery is still functional, which would generate beneficial effects. We obtained support for this concept in the subsequent study with mTORi INK when administered to TRGL/Q175 at 6 months of age for three weeks.

### Autophagy as a primary mechanism for INK-enhanced clearance of small mHTT species

One of the key findings from this study is the reduction of mHTT-IR in neurons detected by immunolabeling after the 3-week oral treatment with mTORi INK (Fig. 7). This beneficial outcome from the drug treatment is interpreted as a primary result of the manipulation/stimulation of autophagy by the compound, leading to enhanced autophagy clearance of mHTT species including smaller, presumably soluble, species and larger aggregates such as the IBs. This interpretation is supported by the following considerations and evidence. First, in theory, the compound used for the treatment is a mTORi so is expected to enhance autophagy [8, 29]. Second, target engagement is achieved as clearly reflected by the diminished levels of mTOR-mediated phosphorylation on mTOR itself (S2481), ULK1 (S757) and p-S6 (S240/S244), along with changes in proteins downstream of mTOR, such as p-ATG14 (S29) and LC3. Third, hue-angle based confocal image analysis of AL/LY subtypes reveals a reduced AL number and an increased LY number in INK-treated TRGL/Q175 compared to Veh-treated TRGL/Q175. The findings from this analysis of AL/LY subtypes highlights the advantages of crossing TRGL with Q175 mice to evaluate autophagy modulators on the ALP, consistent with a notion of developing better tools capable of investigating autophagy flux *in vivo* [50, 74].

Our confocal microscopy analyses revealed mHTT signal in CTSD-positive vesicles, which can be classified as AL based on their LC3 fluorescence, and IEM detected mHTT-silver-enhanced gold particles in AP and AL including lipofuscin granules. Both findings indicate the existence of mHTT species inside the ALP as autophagy substrates. The presence of low levels of mHTT in CTSD-positive AL also implies a constitutive functional degradation event of mHTT in AL (Fig. 2B1, 10-mo-old mouse). This is in contrast to the case in young AD mouse models where early-onset autophagy-stress exhibited severe substrate accumulation (e.g., reflected by strong LC3 signal) due to deficits of degradative functions within AL. [37]. A reduced level in the portion of AV-associated mHTT signals (Fig. 7B2) by mTORi INK further suggests that mHTT degradation has been enhanced after manipulating the ALP. This AV-associated mHTT pool is considered to be corresponding to the mHTT-silver-enhanced gold signal under IEM (Fig. 2B2), i.e., amorphous and existing within the lumen of AV. Such mHTT localization and ultrastructural features are consistent with recent findings that mHTT proteins exogenously added to Q175 brain sections for incubation are recruited to multivesicular bodies/amphisomes, AL and residual bodies (lipofuscin granules), and ultrastructurally localized to non-fibrillar, electron-lucent regions within the lumen of these organelles [82].

### Possible autophagy-related mechanism for the clearance of large mHTT IBs

It is notable that the AV-non-associated form of mHTT (i.e., did not show colocalization with the endogenous tfLC3 signal implying that they might not have a contact with AVs) decrease more obviously after drug-treatment, along with the reductions in p62 and Ub signals (Fig. 7B3). These mHTT species are mainly larger aggresomes/IBs, including those within the nuclei. Based on our mHTT-IEM observation (Fig. 1B), the IBs in Q175 brain are fibrillar aggregates in the nucleus and cytoplasm, exhibiting a homogeneous feature in shape (i.e., round/oval cotton-ball) and in content (i.e., short fine fibrils), similar to the nuclear IBs we found in human HD brains [9], whose cytoplasmic and neuritic IBs, however, also exhibit more heterogeneous compositions such as a mixture of fine fibrils with AVs, fingerprint-like structures or bundles of microtubules. In the current study, we did not observe that a whole IB is contained within an AP or AL in our EM examination, raising a question of how these larger aggregates could be degraded by the autophagy machinery after an autophagy activation by mTOR inhibition. This issue is informed by findings that the mHTT species are dynamic and the aggregates can form as different phases (liquid-like or solid-like), making it possible for mHTT species to move out or into IBs [2, 41, 56, 60] and to shuttle between the nucleus and the cytoplasm [4, 79]. Thus, it is speculated that under the autophagy activated condition induced by mTOR inhibition, the enhanced clearance of mHTT in the ALP pool would promote movements of mHTT species from the IB forms, leading to a reduction on the overall mHTT load.

### Possible additional mechanisms for the reduction of mHTT species

Even if INK is a mTORi mainly targeting autophagy, we cannot completely exclude additional contributions from the UPS for the observed reduction of mHTT species given the crosstalk between the two proteolytic systems. It is well accepted that autophagy plays a more important role in the clearance of larger protein aggregates such as aggresomes/IBs [65]. For example, autophagy is involved in the degradation of p62 bodies [10] and autophagy markers are colocalized with cytoplasmic mHTT IBs [23]. However, there is also evidence for degradation of polyQ aggregates in the nuclei [24], which contain p62 condensates as a hub for UPS-mediated nuclear protein turnover [19]. We observed a high level of colocalization of mHTT with p62 and Ub in aggregates at both intranuclear and extranuclear locations (Fig. 2A), which is presumably more related to the UPS hub and autophagy, respectively. It is speculated that the reduction of mHTT in the cytoplasm due to the stimulation of autophagy may partially relieve the burden of the proteasome in both the cytoplasm and the nucleus so that the nuclear proteasome operates more effectively. One of the observations reported here which may support the above speculation is the reductions of AV-non-associated forms of mHTT/p62/Ub (Fig. 7B3), given that some of these aggregates should exist within the nucleus and therefore their reduced levels (in the nucleus) may reflect increased intranuclear UPS activity, besides the other possibility that they may travel from the nucleus to the cytosol for clearance (by autophagy) as discussed above.

Lowering mHTT via interfering protein production (e.g., through RNAi, antisense oligonucleotides) has been an attractive strategy in HD therapeutic development [34, 69]. Given that mTOR regulates multiple cellular pathways including protein synthesis [42, 77], the inhibition of mTOR as what done in the present study would potentially affect protein synthesis as well. Thus, while our results of decreases in mHTT signals (Fig. 7) can be interpreted as a result of autophagy-mediated clearance of mHTT, a possibility cannot be excluded that mTOR inhibition may result in a reduction in HTT production which may also contribute to the observed results – future studies should determine how significant of such a contribution is.

In summary, Q175 models, as detected in the current study, develop ALP alterations late in the disease progression. Although milder than what is seen in late-stage HD human brain [9], there are similarities in ALP pathobiology between human HD brains and Q175 brains. These include late-onset nature, mHTT as an ALP substrate, increased AL/pa-AL, mouse IB ultrastructural feature similar to the common fine fibril type of human IBs. The late-onset mild ALP pathology in Q175 is different from the severe AV accumulation in mouse models of AD as mentioned above [37]. Together, the mild and late-onset nature of ALP alterations in Q175 provides a basis for manipulating autophagy to be a promising therapeutic strategy if the manipulations (e.g., stimulating autophagy with mTORi as in this study) are applied when the autophagy machinery is largely undamaged. This has been validated to be successful in this study as demonstrated by the

INK-induced reduction of mHTT species along with other beneficial effects, consistent with findings from similar studies and supporting the general notion of lowering mHTT as a therapeutic strategy for HD [6, 14, 59]. However, it should be noted that the current study is an experimental therapeutical attempt in a mouse model which, consistent with previous reports [59], is just a proof of concept for manipulating autophagy (e.g., via inhibiting mTOR in the current setting) as a potential therapeutic. The clinical implications from such studies require further rigorous verifications where early diagnosis and earlier interventions would be critical factors to be considered.

## Declarations

## Acknowledgments

We are grateful to Dr. T Yoshimori (Osaka University, Japan) for the mRFP-eEGFP-LC3 construct used in the TRGL mice.

## Funding

This work was supported by the CHDI Foundation (R.A.N.) and the National Institute of Aging (P01 AG017617 to R.A.N.).

## Author Contributions

Conceptualization and design: RAN, DSY, DMM; Investigation, data collection and analysis: PS, DSY, CNG, JP, CH, VK; Writing - original draft preparation: DSY; Writing - review and editing: DSY, RAN, DMM, PS, CNG, MR; Funding acquisition: RAN; Supervision: DSY, RAN.

## Competing Interest

All authors declare no conflict of interests.

## Ethics Approval

The animal procedures conducted in this study were approved by the Institutional Animal Care and Use Committee at the NKI.

## Abbreviations

AD: Alzheimer’s disease
AL: autolysosomes
ALP: autophagy-lysosomal pathway
AP: autophagosomes
ATG: autophagy-related protein
AV: Autophagic vacuoles
BBB: blood-brain barrier
CTSB: cathepsin B
CTSD: cathepsin D
DARPP-32: dopamine- and cAMP-regulated phosphoprotein, 32 kDa
HD: Huntington’s disease
HTT: huntingtin protein
IB: inclusion bodies
IEM: immuno-gold electron microscopy
IF: immunofluorescence
IHC: immunohistochemistry
INK: mTOR inhibitor INK-128
IR: immunoreactivity
LAMP1: lysosomal-associated membrane protein 1
LC3: microtubule-associated protein 1 light chain 3
LM: light microscopy
LY: lysosomes
mTOR: mechanistic target of rapamycin kinase
mTORi: mTOR inhibitor
mHTT: mutant huntingtin protein
NII: neuronal intranuclear inclusions
pa-AL: poorly acidified AL
Q175: B6J.zQ175 knock-in mice
SQSTM1/p62: sequestosome 1
STR: striatum
tfLC3: tandem fluorescent mRFP-eGFP-LC3
TRGL: Thy-1-RFP-GFP-LC3
Ub: ubiquitin
ULK1: unc-51 like autophagy activating kinase 1
UPS: ubiquitin-proteasome system
WB: western blotting
WT: wild type.

---

**Fig. S1 (related to Fig. 2).**
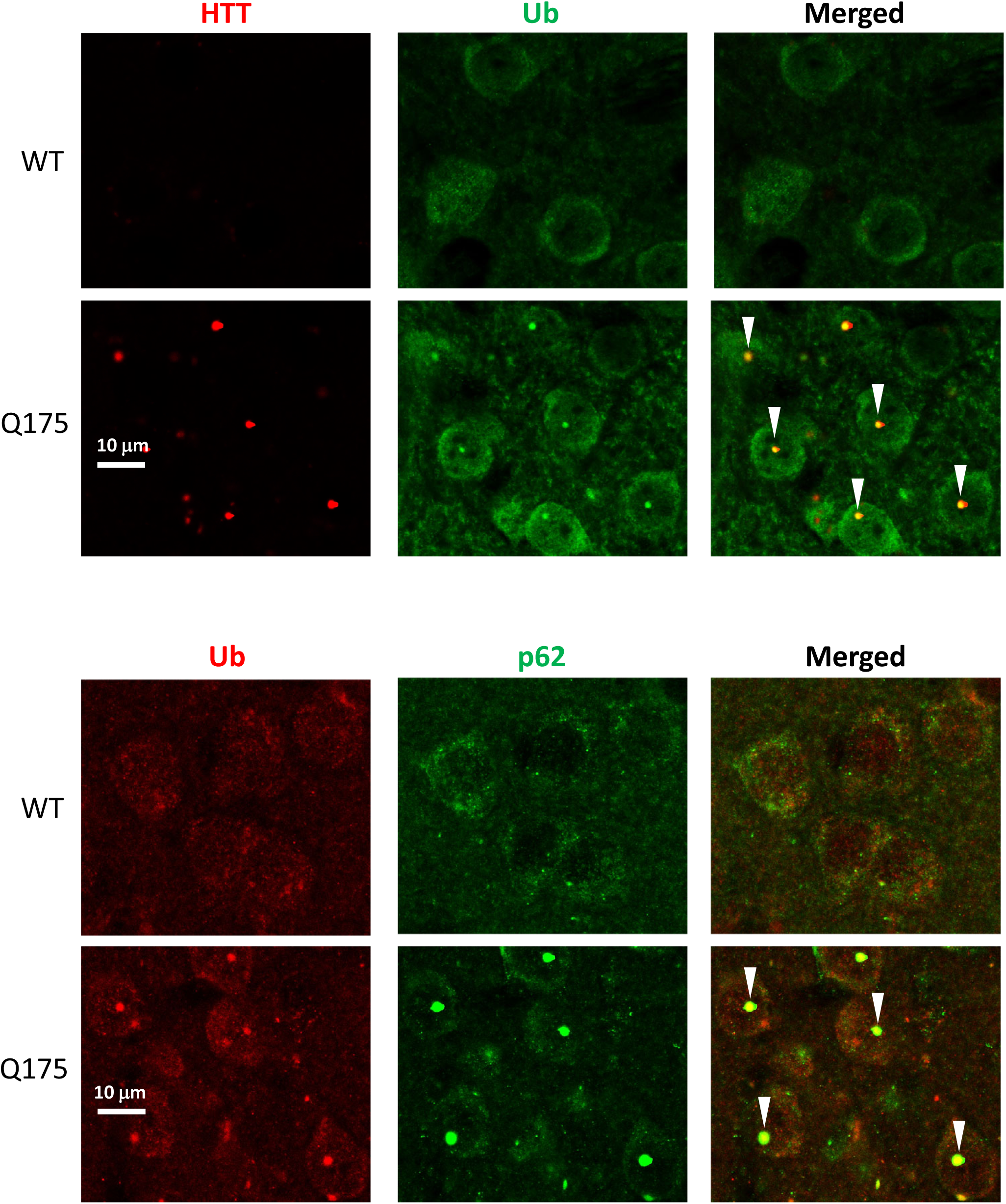
Colocalization of HTT/Ub and Ub/p62 in IBs in the STR. Brain sections from 10-mo-old mice were processed for double IF with antibodies against HTT (antibody MW8; red) and pan-Ub (green), or pan-Ub (red) and p62 (green), and confocal images from the striatum are shown. Arrowheads depict IBs showing colocalization signals.

**Fig. S2 (related to Fig. 3).**
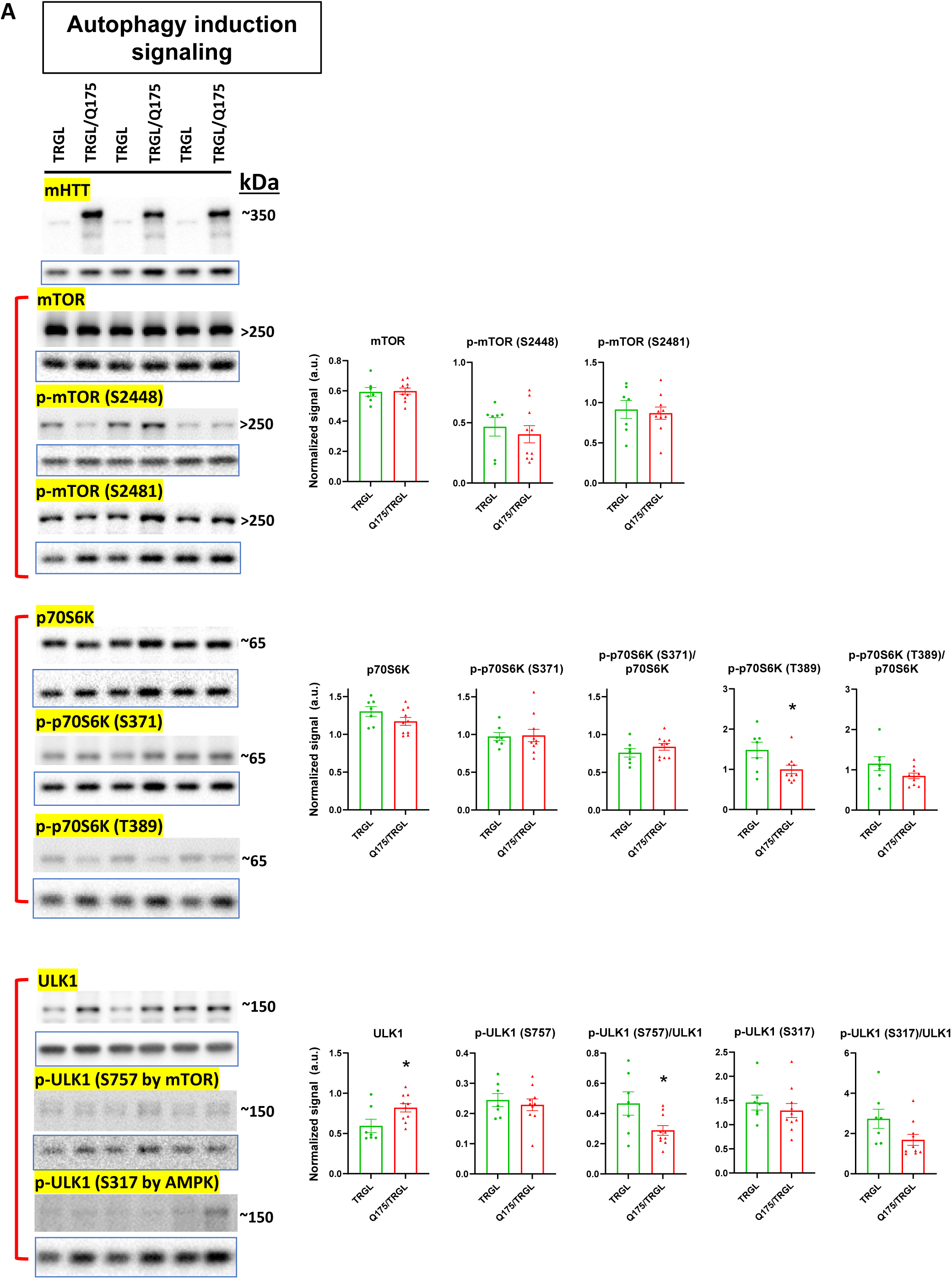
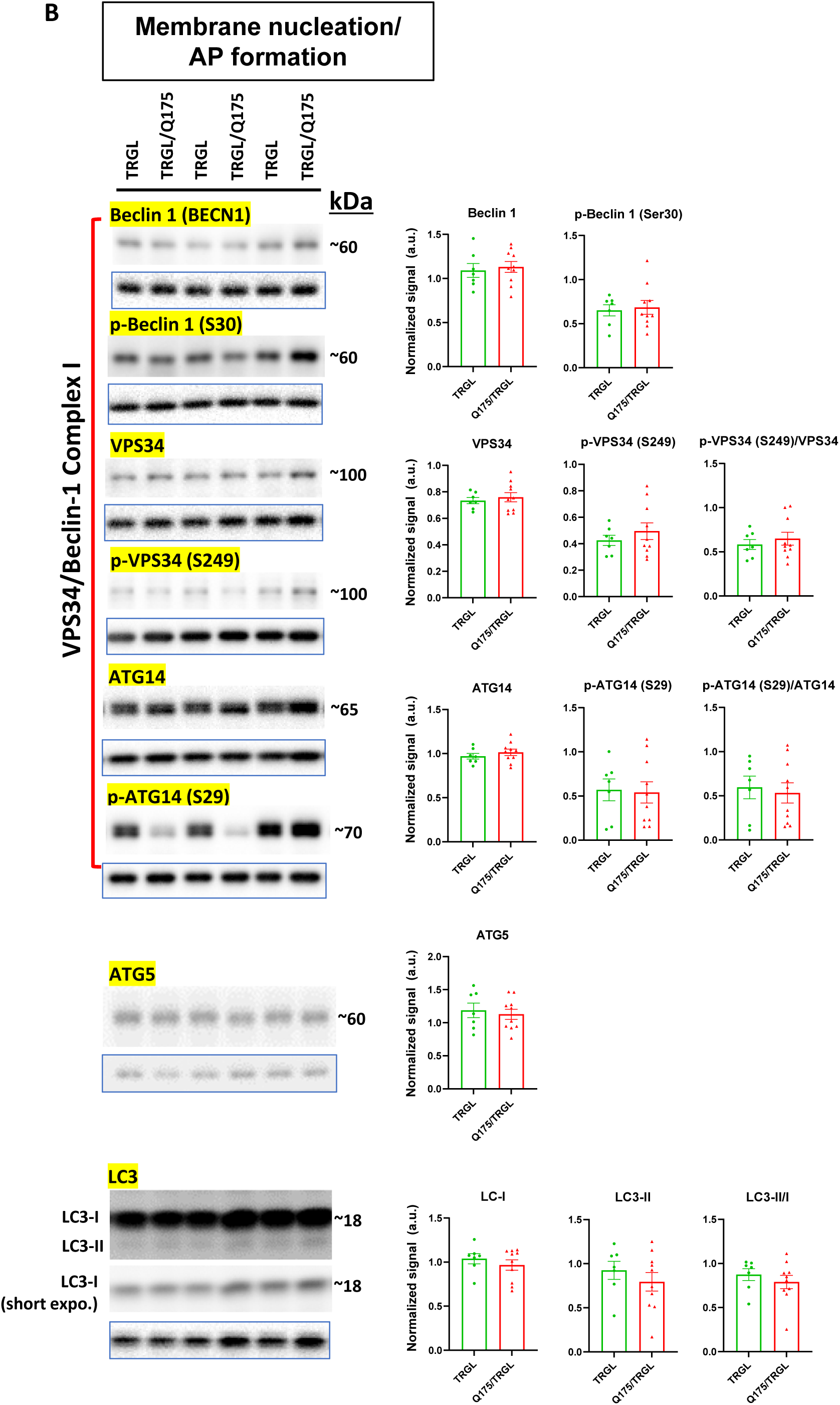
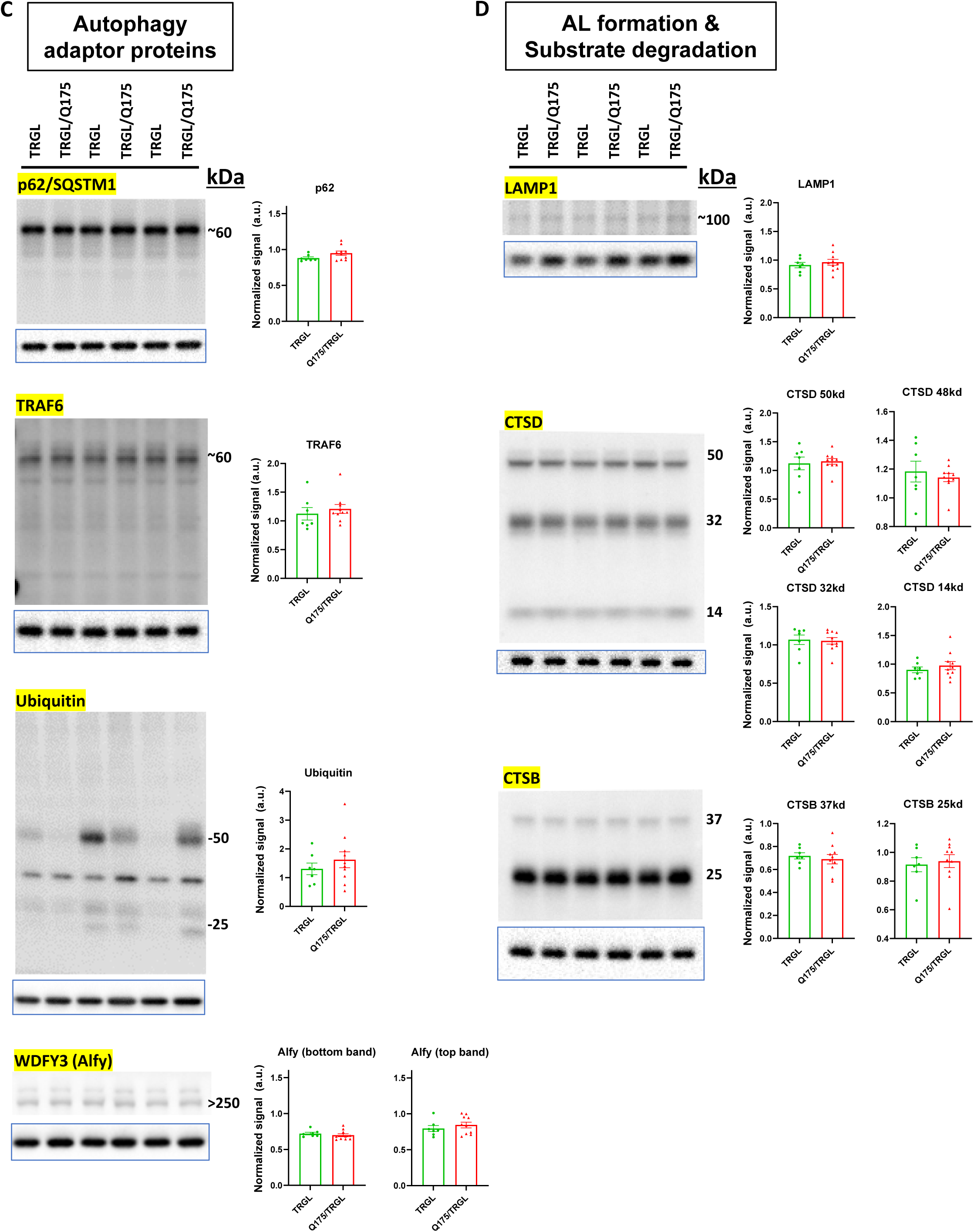
Molecules involved in various stages of the ALP are largely unchanged in 17-mo-old TRGL/Q175. Equal amounts of proteins from brain homogenates of 17-mo-old TRGL and TRGL/Q175 were subjected to SDS-PAGE and processed for WB with antibodies directed against a number of interested marker proteins in the ALP, grouped according to the functions of the proteins in the autophagy process: autophagy induction signaling (A), membrane nucleation/AP formation (B), autophagy adaptor proteins (C) and AL formation/substrate degradation (e.g., lysosomal hydrolases or structural components)(D). The blot of the loading control protein GAPDH is boxed and placed under each protein of interest. Images were collected by a digital gel imager (Syngene G:Box XX9). Densitometry was performed with Image J and the result for each protein of interest was normalized by its corresponding GAPDH blot and presented in the bar graph. Values are the Mean +/- SEM for each group (n = 7 TRGL and 10 TRGL/Q175). Significant differences between the 2 groups were analyzed by unpaired, two-tailed t-test. * P < 0.05, ** P < 0.01, *** P < 0.001.

**Fig. S3 (related to Fig. 5).**
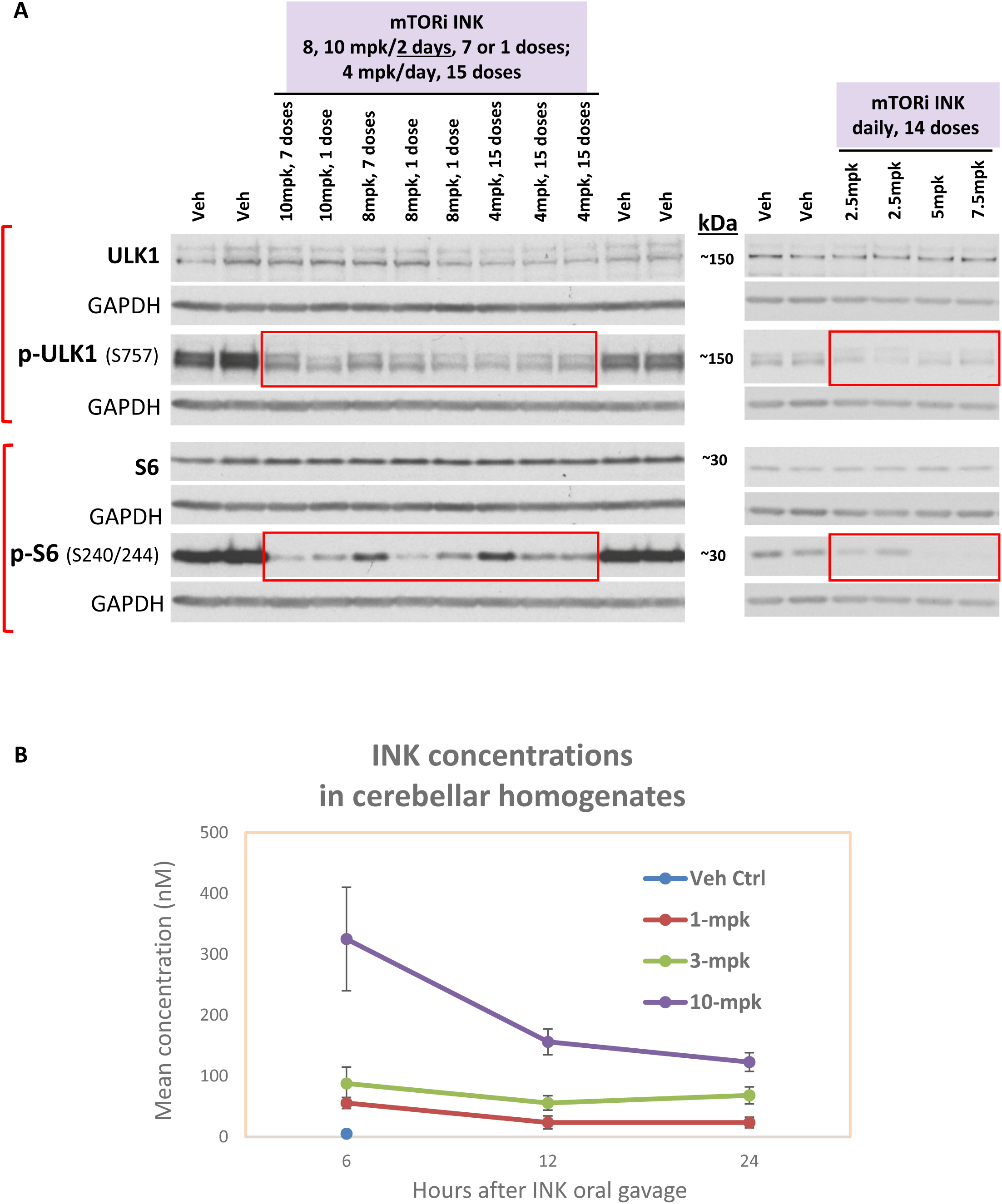
Dosing tests demonstrate mTORi INK BBB penetration and target engagement even at low dosages. ***(A) Dosing tests for INK.*** INK or the vehicle was administered by oral gavage. Two dosing tests were done in 6-mo-old WT mice, one with 10, 8 mg/kg (mpk) every other day or 4-mpk daily for 2 weeks and the other with 7.5, 5, 2.5-mpk daily for 2 weeks. Equal amounts of proteins from brain homogenates of mice receiving vehicle (labeled as “Veh”) or INK were subjected to SDS-PAGE and processed for WB with antibodies directed against marker proteins representing target engagement of INK, such as p-ULK1 (S757) and p-S6 – the drug effects are reflected by the changes within the boxed areas compared to the signals of Veh outside the boxed areas. ***(B) INK was detected in the brains of mice receiving a single dose via oral gavage.*** After 6, 12 or 24 hrs post INK administration to 6-mo-old WT mice, INK was detected in cerebellum of mice treated with 1, 3 or 10-mpk INK. Total 60 mice: n = 6 mice/dose/time point, where the Vehicle control group was just tested once at the 6-hr time point.

## Notes

### Competing Interest Statement

The authors have declared no competing interest.

### Summary of Updates

In response to the reviewers' comments from the peer review process by eLife, this version contains minor modifications in Fig. 5A, Fig. 6A and Discussion.

